# Single-cell transcription mapping of murine and human mammary organoids responses to female hormones

**DOI:** 10.1101/2023.09.28.559971

**Authors:** Jenelys Ruiz Ortiz, Steven M. Lewis, Michael F. Ciccone, Deeptiman Chatterjee, Samantha Henry, Adam Siepel, Camila O. dos Santos

## Abstract

During female adolescence and pregnancy, rising levels of hormones result in a cyclic source of signals that control the development of mammary tissue. While such alterations are well understood from a whole-gland perspective, the alterations that such hormones bring to organoid cultures derived from mammary glands have yet to be fully mapped. This is of special importance given that organoids are considered suitable systems to understand cross species breast development. Here we utilized single-cell transcriptional profiling to delineate responses of murine and human normal breast organoid systems to female hormones across evolutionary distinct species. Collectively, our study represents a molecular atlas of epithelial dynamics in response to estrogen and pregnancy hormones.

## Introduction

During adolescence, a surge in hormones estrogen (E2) and progesterone (P4) transform the rudimentary mammary epithelium developed during embryogenesis into a complex epithelial hierarchy, marked by lineage defined cells with distinct functions (Naccarato et al., 2000; Richert et al., 2000). Yet, the most drastic postnatal developmental stage of the mammary gland occurs during pregnancy, due to the interplay of elevated levels of E2, P4 and prolactin (PRL), which collectively induce the maturation of the mammary gland into a milk secretory organ (Gallego et al., 2001; Mueller et al., 2002; Ruan et al., 2005; Slepicka et al., 2021).

Although mouse models have been extensively used to assess the heterogeneity of the mammary gland and its associated developmental processes, organoid systems are emerging as an attractive model system that allows for the 3D culturing of mammary fragments under conditions that resemble the *in vivo* environment (Nguyen-Ngoc et al., 2015). Previous studies have shown that human normal mammary-derived organoids are able to recapitulate MEC lineage diversity *ex vivo* (Gray et al., 2022; Rosenbluth et al., 2020), thus representing a scalable and easily applicable model to studying the role of soluble mediators in modifying MEC function. Additionally, when grown with combinations of diverse hormone and factors, mammary-derived organoid cultures activate molecular dynamics that resemble those present during pregnancy, lactation and involution, making this system suitable to define the effects of different hormones on MECs at determined concentrations and time points (Ciccone et al., 2020; Stewart et al., 2021; Sumbal et al., 2020). As organoids gain traction as a model system for the study of the mammary gland in developmental biology and in cancer, a deeper characterization of this system, for mice and human cells, and the extent to which they fully recapitulate *in vivo* biology is needed (Lewis et al., 2022).

Here, we set out to define the molecular and cellular changes brought upon by supplementation with a pregnancy-hormone cocktail, in both murine and human organoids using single-cell RNA sequencing (scRNA-seq). We determined that murine organoid models capture the heterogeneity of intact mammary epithelial tissue, an analysis that revealed the existence of phenotypes exclusive to *ex vivo* cultures, marked by pathways associated with less differentiated cellular states. We also characterized the response of murine organoids with different doses of estrogen, as a way to integrate important mammary developmental and maintenance signals to *ex vivo* derived systems, an approach that identified molecular dynamics of hormone responsive and sensing cellular states. The single-cell characterization of mammary organoids systems grown with pregnancy-associated hormones allowed for the identification of additional cellular responses beyond those induced by estrogen. Here, the utilization of data prediction approaches allowed for the comparison of pregnancy-induced organoid states to those observed during pregnancy in mice. This analysis further illustrated the pregnancy mimicking potential of *ex vivo* systems, further supporting its potential as a model system to understand hormone driven mammary developmental stages.

An important advantage of utilizing 3D organoid cultures to investigate normal mammary gland development rely on the opportunity to study such process in tissues where *in vivo* studies are less accessible (Finot et al., 2021; Mackenzie et al., 1982; Ogorevc & Dovč, 2015; Sun et al., 2005). Therefore, we further explored the robustness of mammary derived organoids, to study pregnancy-induced development of human MECs. Here we utilized organoid systems already previously described to represent the tissue cellular heterogeneity of freshly isolated breast tissue (Bhatia et al., 2022). Initial characterization of these systems indicated the expression of lineage defining markers across all organoid cellular states, indicating expression infidelity induced by the culturing system. We therefore derived a set of *de novo* markers of cellular identities of organoid grown human MECS, by combining differentially expressed genes across untreated and pregnancy hormones treated systems, thus capturing alterations driven by signals that influence cellular behaviors.

By employing analytical cross species approaches, we also demonstrated distinct and evolutionarily conserved cellular and molecular dynamics of pregnancy hormone responses. Altogether, our efforts have generated a single-cell map of murine and human MEC-derived organoids undergoing hormone response *ex vivo*. This resource has the potential to pave the way for future studies exploring 3D systems to model mammary tissue development.

## Methods

### Animals

Nulliparous female C57BL/6 mice were purchased from Jackson Laboratory. All animals were housed in a 12-hour light-dark cycle with controlled temperature and humidity at 72°F and 40-60%, respectively, with access to dry food and water ad libitum. All animal experiments were performed in accordance with the CSHL Institutional Animal Care and Use Committee.

### Murine Organoid Derivation and Culture

Mammary-derived organoid cultures were cultured as previously described (Ciccone et al., 2020), within Matrigel (Corning, CATALOG INFO) domes, submerged in Advanced DMEM/F12+++ media supplemented with 1X ITS (Insulin/Transferrin/Sodium Selenite, Gibco #41400-045) and FGF-2 at 5 nm (PeproTech, Cat# 450-33): essential medium. Organoid culture media was changed every 2 days. FGF-2 was then withdrawn from the organoid cultures for 24 hours after which the treatment regimen was initiated. Organoid conditions with “low” levels of estrogen were grown with media supplemented with 33.3 ng/mL of 17-β-Estradiol (Sigma #E2758), and those with “high” levels of estrogen were grown in the presence of 66.6 ng/mL of 17-β-Estradiol. Mouse organoid conditions to mimic pregnancy were cultured with media supplemented with 66.6 ng/mL of 17β-Estradiol, 200 ng/mL of progesterone (Sigma #P8783) and 200 ng/mL of prolactin (Sigma #L4021). In all conditions, hormone treatment was carried out for 48 hours. For the preparation of scRNAseq, organoid cultures were dissociated with 500 µL of Cell Recovery Solution (Corning® # 354253) for 30 minutes, followed by incubation with 500 µL of cold Tryp-LE (Thermo Fisher Scientific #12604-013) at 37 °C for 10 minutes. Dissociated organoids were resuspended with 1 mL media, transferred to a 15 mL BSA pre-coated Falcon tube, and spun at 300 G for 5 minutes. Dissociated organoid cells were then resuspended in 1 mL of media and submitted for library preparation and sequencing.

### Human Organoids

Established patient-derived normal breast organoid cultures (Bhatia et al., 2022) were cultured as previously described, within Matrigel (Corning) domes, submerged in media containing 10% R-Spondin1 conditioned medium, 5 nmol/L Neuregulin 1 (Peprotech, 100–03), 5 ng/mL FGF7 (Peprotech, 100–19), 20 ng/mL FGF10 (Peprotech, 100–26), 5 ng/mL EGF (Peprotech, AF-100–15), 100 ng/mL Noggin (Peprotech, 120–10C), 500 nmol/L A83–01 (Tocris, 2939), 5 μmol/L Y-27632 (Abmole, Y-27632), 500 nmol/L SB202190 (Sigma, S7067), 1× B27 supplement (Gibco, 17504–44), 1.25 mmol/L N-acetylcysteine (Sigma, A9165), 5 mmol/L nicotinamide (Sigma, N0636), and 50 μg/mL Primocin (Invitrogen, ant-pm-1) in ADF+++. Organoid culture media was changed every 3 days, and organoids were passed every 5-8 days to avoid confluency. Human MEC derived organoids were treated with pregnancy hormone concentrations same to those utilized for the growth of murine organoids (66.6 ng/mL of 17β-Estradiol, 200 ng/mL of progesterone (Sigma #P8783) and 200 ng/mL of prolactin (Sigma #L4021)). We confirmed with qPCR analyses that these grow conditions induced the expression of casein genes, and utilized such analysis to define the collection time points for scRNAseq (untreated cultures, and 10 and 21 after supplementation of media with pregnancy hormones). Cultured human organoids were processed similarly to mouse organoids prior submission for library preparation and sequencing, with organoids being dissociated with 500 µL of Cell Recovery Solution (Corning® # 354253) for 30 minutes, followed by incubation with 500 µL of cold Tryp-LE (Thermo Fisher Scientific #12604-013) at 37 °C for 10 minutes. The dissociated human organoids were likewise resuspended with 1 mL media, transferred to a 15 mL BSA pre-coated Falcon tube, spun at 300 G for 5 minutes, resuspended in 1 mL of media and submitted for library preparation and sequencing.

### qPCR analysis

Organoid MECs were homogenized in Trizol (Thermo Fisher Scientific) for RNA extraction. Double stranded cDNA was synthesized from purified total RNA using SuperScript III Reverse Transcriptase (Thermo Scientific). QuantStudio 6 real time PCR system, Software v1.3 (Thermo Fisher) and quantification results were analyzed using the delta delta CT method. The relative mRNA expression of the target gene was determined using the ΔΔCt method and normalized against β-ACTIN mRNA levels. Comparing *CSN2* and *CSN3* expression at 0 vs 21 days of EPP treatment using the Mann Whitney test yielded a significant difference in expression for *CSN2* at 21 days (p value = 0.0238), while *CSN3* resulted in a non-significant difference in expression (p value = >0.9999).

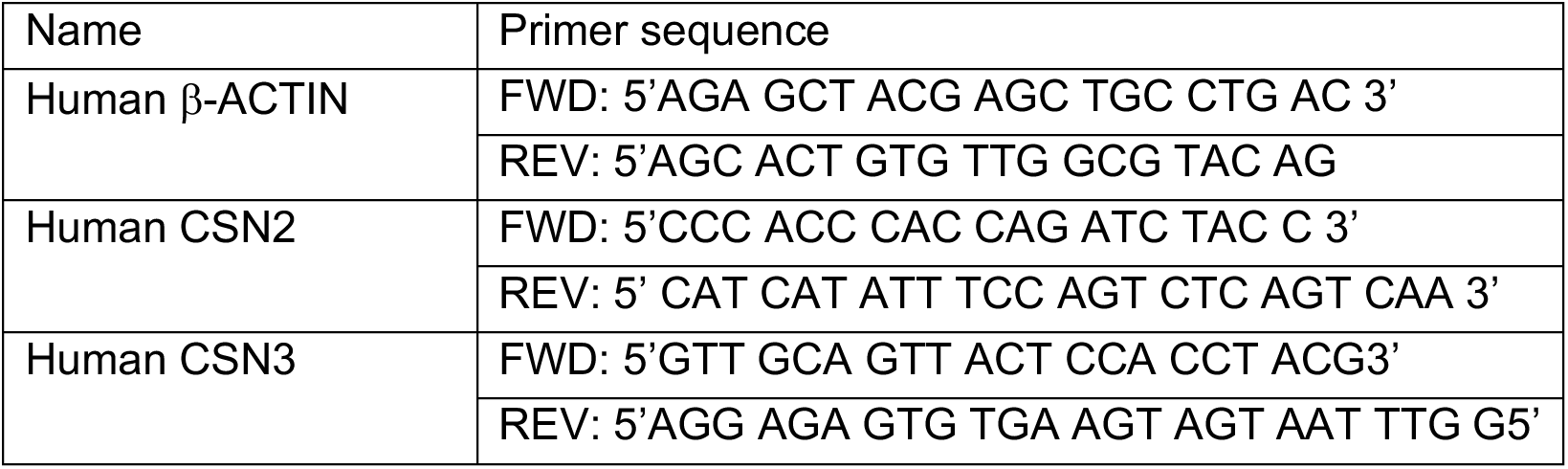

### scRNAseq library preparation and data analysis

Libraries were prepared with the 10X Chromium platform for single-cell libraries. The libraries were run with 3’ chemistry single end sequencing and indexing using the Illumina NextSeq 550 high output platform. Libraries from mouse samples were aligned to the mm10 genome using CellRanger v3, and human libraries were aligned to the GRCh38-2020 genome using CellRanger v6. All further data processing and analysis was completed in the Seurat package in R version 4.0.0. Initial quality control involved removing any cells with mitochondrial RNA expression over 15%, removing clusters with high ribosomal RNA expression and removing clusters with >5,000 and <200 features. For batch effect correction and normalization, anchors were discovered between the datasets using the FindIntegrationAnchors() function before integrating with the IntegrateData() function. Throughout the analysis and re-clustering, repeated quality control through evaluation of clusters with a large proportion of cells expressing low features or high mitochondrial RNA content were removed. This ensured the removal of low-quality clusters at each stage of the processing and analysis. Uniform manifold approximation and projection (UMAP) clustering using a shared nearest neighbor graph (SNN) was performed. The resolution of each clustering step with the help of Clustree (Zappia & Oshlack, 2018), and all of the analysis presented here were run with a resolution of 0.3, with the exception to data analysis shown on Figure 3E, which due to the large number of samples, was performed with a resolution of 0.2. Differences in cell numbers between datasets were analyzed with the Propeller package, which uses a robust and flexible method that leverages biological replication to find statistically significant differences in cell type proportions between groups (Phipson et al., 2022). Regulon analysis for each culturing condition and species was performed using SCENIC version 1.2.0 in R (Aibar et al., 2017).

Identity assignment of epithelial cell clusters was performed using module scores based on known lineage markers (Henry et al., 2021) (**Table 1**), and/or top differentially expressed genes, assigned to each cluster in each Seurat object. To evaluate differentially expressed genes (DEGs) within our data, we utilized the FindMarkers() function, which completes a Wilcoxon rank-sum test to identify DEGs between clusters. For visualizing DEGs and particular genes of interest within the data, we utilized the following functions: DotPlot(), FeaturePlot(), VlnPlot() and HeatMap(). For a dendrogram analysis of the relative relatedness of the clusters, we utilized the BuildClusterTree() function using default parameters. Ternary plots were generated from resulting module scores for broad MEC lineage markers using the ternary plot() function from the vcd package. The naming of cell types is in accordance with Human Breast Cell Atlas (HBCA) discussions.

**Table 1.** General lineage markers for murine MECs.

| Mouse MEC lineage | Gene markers |
| --- | --- |
| Luminal Hormone Sensing (LHS) | Epcam, Krt8, Krt18, Prlr, Prrg2, Ak3, Cdk19, Fxyd2, Areg, Stc2, Prom1, Esr1, Pgr, Cdo1, Gstm2, Wnt5, Cxcl15, Ly6a, Tspan9, Gltp, Cd14, Ppme1, Adck5, Dusp4, Tph1, Notch3, Itpripl1 Calca, Armcx2, Cited1, Rcan1, Pak6, Pir, Fgb, Fam83g, Il6ra, Itpripl2, Ptbp2 |
| Luminal Adaptive Secretory Precursor (LASP) | Epcam, Krt8, Krt18, Col9a1, Il1rn, Itga2, Csn1s1, Car2, Csn2, Bptf, Lalba, Kit, Armcx2, Trf, Cxcl1, Ndst1, Ezh2, Ap1g1, Areg, Spp, Sfxn3, Cd14, Snx27, Mfsd5, S100a8, Lbp,, Gjb2, Notch3, Il6ra, Kctd20, Erf, Ptbp2, Ireb2, Csn3, Aldh1a3, Ceacam1, Csf3, Bmpr2, Egln3, Il1m, Lgals1, Stmn1, Tgfb3, Mki67, Lockd, H2afz, Ube2c, Prrg2, Ak3,, Cdk19, Mdk,, Ly6a, Krt14, Top2, Tagln, Cenpa, Fam83g, Rangrf, Ppme1, Hmgb2, Itpripl2, Setd7, Cwc22, Parp1, Sms, Sp110, Cenpa, Fam83g, Rangrf, Ppme1, Hmgb2, Itpripl2, Setd7, Cwc22, Parp1, Sms, Sp110, Cxcr4 |
| Basal Myoepithelial (BMyo) | Epcam, Lgals1, Bptf, Krt17, Ppic, Mdk, Krt14, Krt5, Acta2, Mgp, Lmod, Lhfp, Cxcl14, Serpina3n, Cnn1, Vcam1, Nrg1, Col7a1, Nexnll17b, Mylk, Sparc, Lgr5, Jag1, Scn7a, Trp63, Lbp, Tagln, Bmpr2, Fgf1, Lipg, Arcld4, Mme, Mmp2, Igfbp3, Oxtr |

For data presented on Figure 1A, organoids derived from MECs of 3 never pregnant, nulliparous female mice were utilized on the generation of scRNA-seq libraries, using the 10X Chromium platform, yielding a total 10,508 Mouse Organoid (MO) cells. For data presented on Figure 1C, only epithelial cells (*Epcam*+, *Krt5*+, *Krt14*+, *Krt8*+, and *Krt18*+) were selected from publicly available, intact mammary tissue scRNAseq datasets, resulting in 1,986 cells originating from the Henry et al. data set and 4,025 from Bach et al. (Bach et al., 2017; Henry et al., 2021). After integration with mammary organoids scRNAseq (**Fig. 1C**), a total of 6 Organoid-MECs Integrated with Mouse-MECs (OIM) clusters, composed of 10,502 cells from organoid cultures and 6,011 cells from intact mammary tissue. An initial batch effect correction was performed for the merging of the Henry et al. data set, both Bach et al. samples and our organoid data set to account for the different number of cells in both organoids and intact tissue samples and any technical variability.

**Figure 1.**
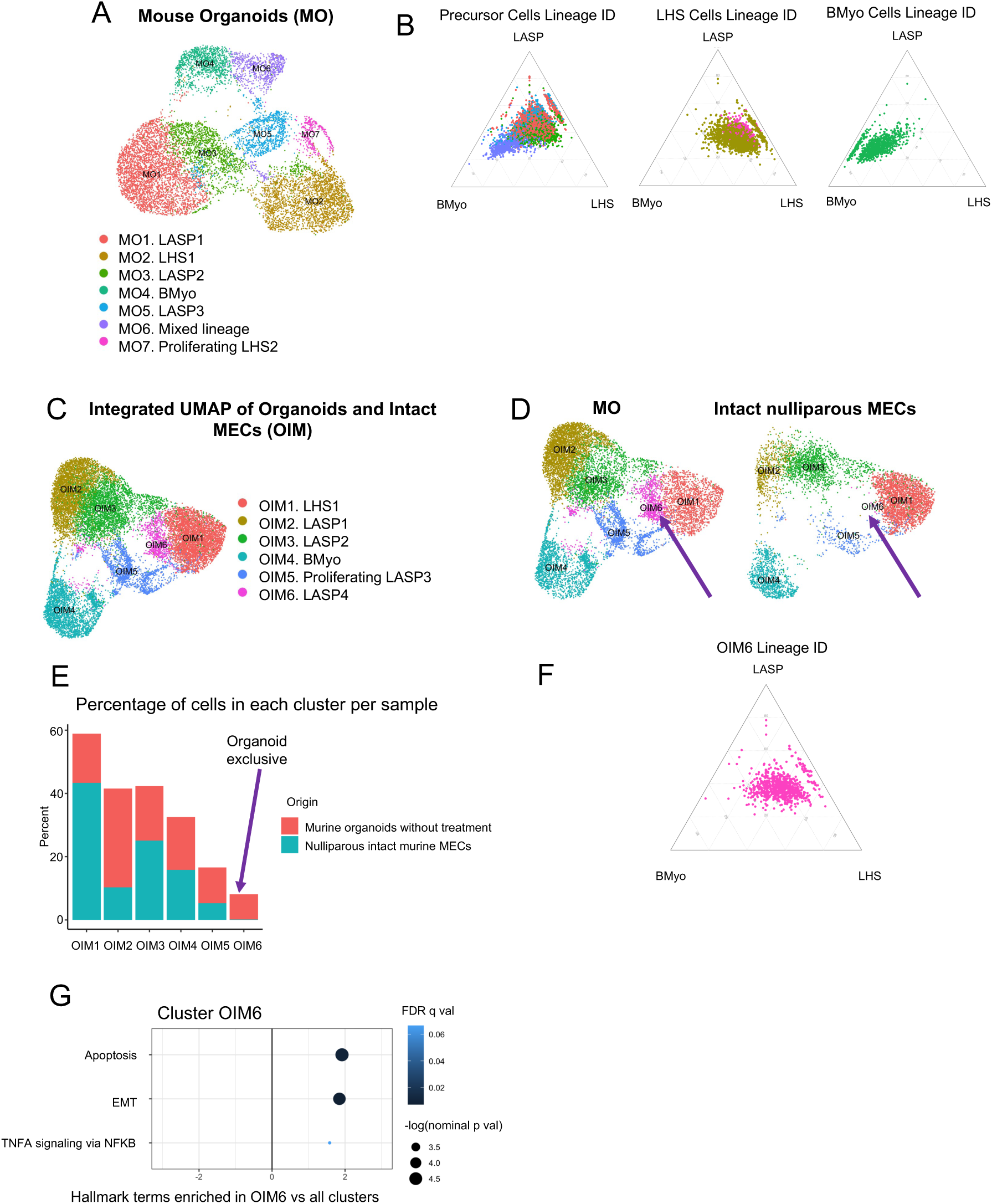
Analysis of mouse organoid MECs scRNA-seq data. (A) Mouse Organoid (MO) clusters and their given identities according to gene expression from previously described MEC markers. (B) Ternary plots showing how each MO cluster scores for general lineage markers (Table 1). MO clusters are organized based on their dendrogram relationships. (C) Integrated analysis of Organoids and Intact MECs (OIM) clusters and their given identities according to gene expression from previously described MEC markers. (D) OIM clusters split by condition (cells originating from organoids or from intact tissue). The purple arrow is highlighting OIM6, a cluster of luminal progenitors that appears to be enriched in organoid cultures. (E) Bar plot showing percentage of cells per condition in each cellular cluster. The purple arrow highlights OIM6, an organoid exclusive cluster. (F) Ternary plot showing how OIM6 cells score for general lineage markers (Table 1). (G) GSEA for hallmark terms enriched in cluster OIM6. Hallmark terms are ordered based on the –log of nominal p-values for each term. Only terms with a nominal p-value (nom p-val) < 0.05 were kept for this analysis, in order to only show significantly enriched terms. The dots are colored based on their false discovery rate (FDR q-value), and the x-axis represents normalized enrichment scores (NES).

For data presented on Figure 2, organoid cultures treated with estrogen concentrations for 48 hours were prepared for scRNA-seq with the 10X Chromium platform. Quality control filtering steps and clustering alongside the untreated murine MEC-derived organoids, yielded a total of 9 clusters containing 31,802 Organoid-MECs with estrogen (OE). From these, 10,508 cells were from untreated samples, 9,695 cells were from samples treated with a low dose of estrogen (33.3 ng/mL), and 11,599 cells were from samples treated with a high dose of estrogen (66.6 ng/mL). Each of the cell cluster identities were determined once more using previously described lineage commitment markers in intact mammary tissue (Henry et al., 2021).

**Figure 2.**
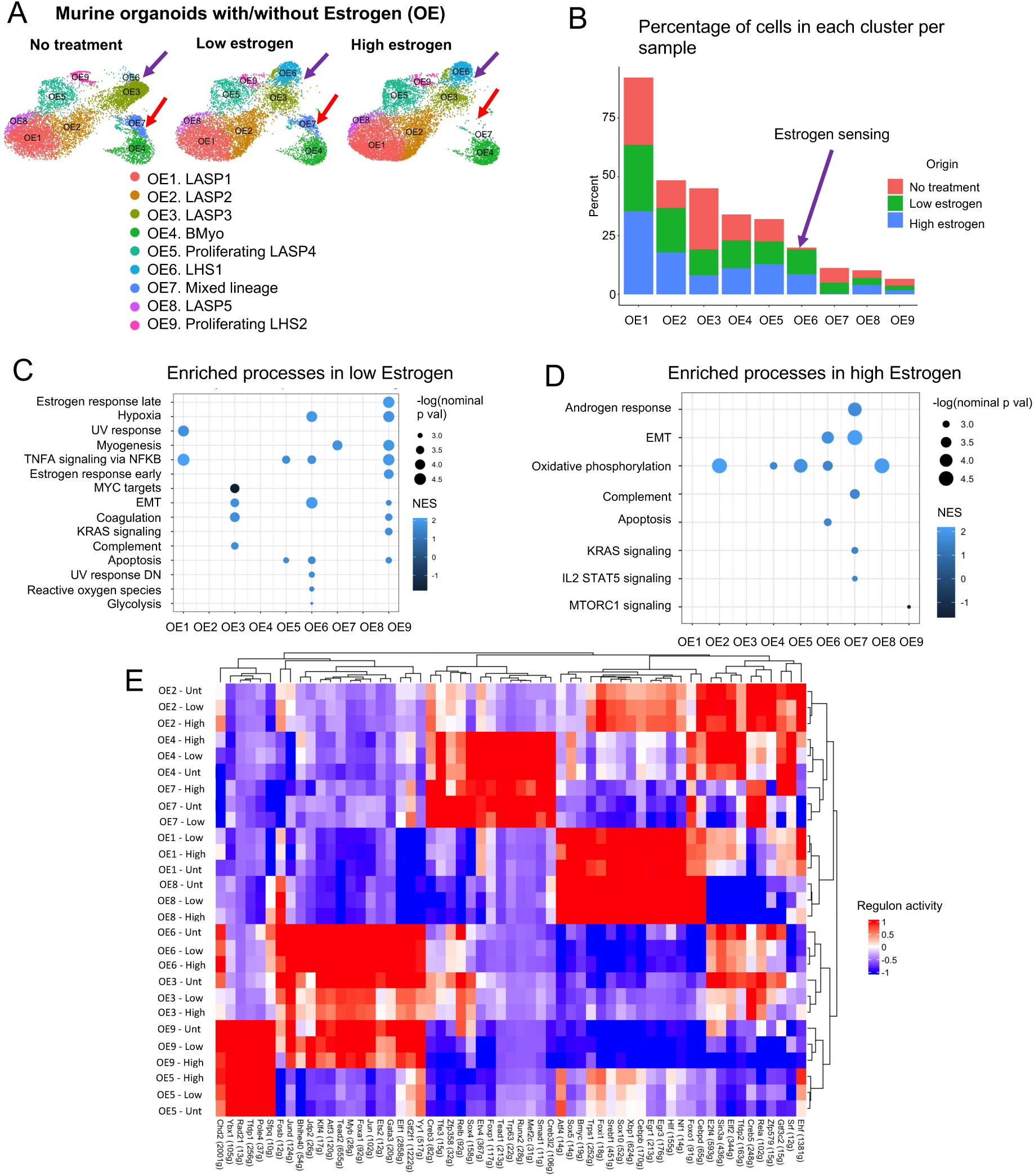
Analysis of scRNA-seq data from mouse organoid MECs with Estrogen treatment. (A) OE clusters split by condition, highlighting Estrogen-specific cluster OE6 (purple arrow) and OE7 (red arrow), which is depleted only at a high Estrogen dose. (B) Bar plot showing percentage of cells per condition in each cellular cluster. The purple arrow highlights OE6, an Estrogen-exclusive cellular cluster. (C-D) GSEA for hallmark terms differentially enriched in each Estrogen treatment condition across clusters. Terms were ordered decreasingly based on their–log(nom p-value). Only terms with nom p-val <0.05 were kept for these analyses. The color of each dot represents the NES for each term. (E) Regulon analysis showing the activities of regulons with the highest RSS per cluster and condition. The activities of each regulon are scaled to represent significant activity in one cluster or condition (red), and no significant activity (blue).

For data presented on Figure 3, quality control steps and clustering of datasets from organoids without treatment, and those treated with estrogen, progesterone and prolactin (EPP), resulted in 10 Organoids with/without EPP (OP) clusters, with a total of 26,971 cells, 10,508 from our untreated samples and 16,463 from our samples treated with EPP. Untreated organoids and those treated with EPP were also merged with publicly available datasets from murine mammary tissue collected at different pregnancy stages (Bach et al., 2017). After QC filtering, we obtained a total of 7 organoids integrated with MECs from a pregnancy (OIP) clusters, with a total of 4,004 cells from nulliparous (NP) MECs, 5,216 MECs from mice during gestation, 8,222 from mice during lactation, 5,607 from mice during involution, 10,497 untreated organoid cells, and 16,449 EPP-treated organoid cells.

**Figure 3.**
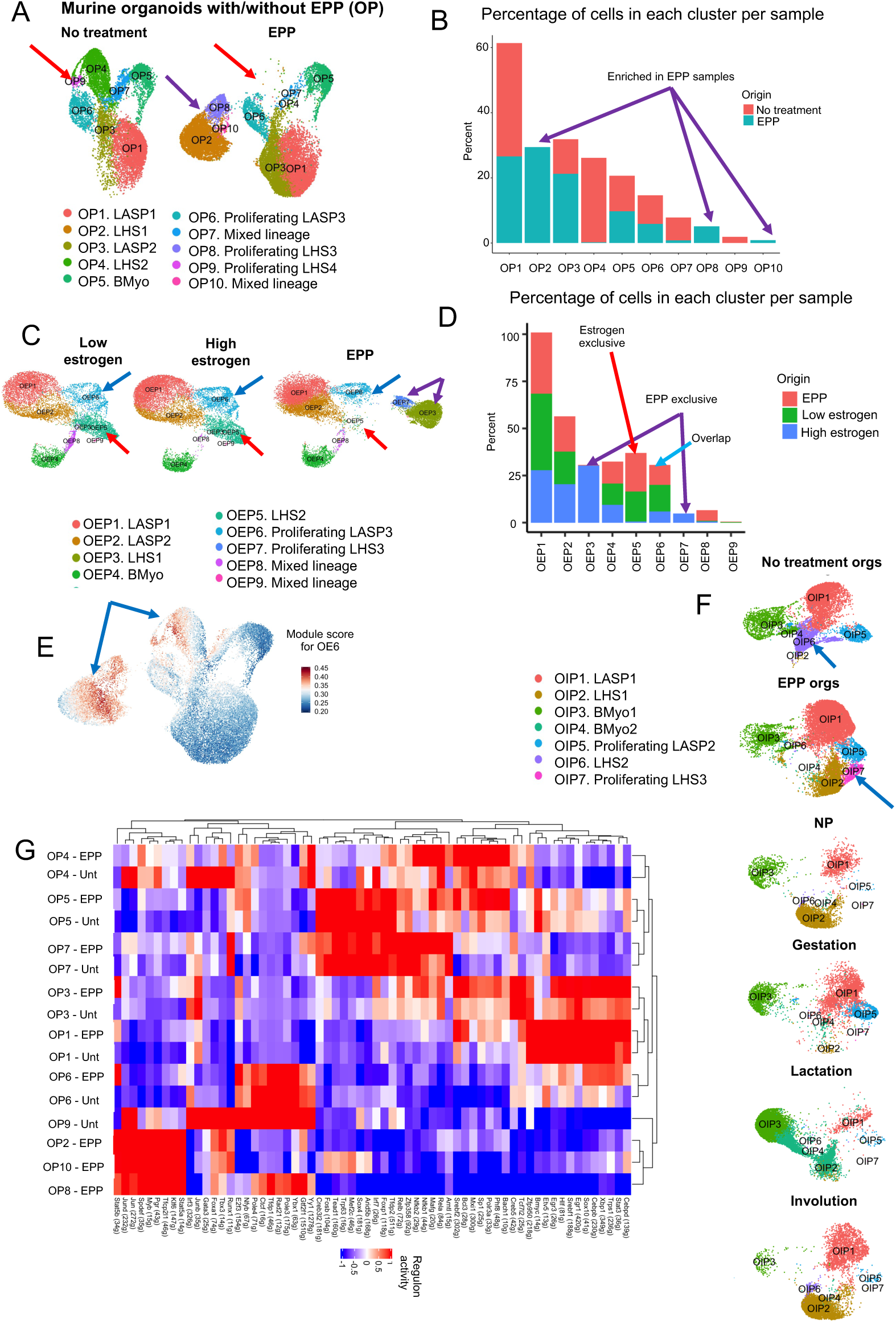
Analysis of scRNA-seq data from murine organoid MECs with EPP treatment. (A) OP clusters split by treatment condition (untreated or EPP treatment). The purple arrow highlights EPP-enriched cellular clusters OP2, OP8 and OP10. The red arrow highlights cellular clusters depleted with EPP treatment, clusters OP4, OP7 and OP9. (B) Bar plot showing percentage of cells per condition in each OP cluster. The purple arrows highlight clusters enriched in EPP samples. (C) Analysis of scRNA-seq data from organoids treated with Estrogen or pregnancy hormones (OEP). The purple arrows highlight EPP-exclusive clusters OEP3 and OEP7, red arrows highlight estrogen-exclusive cluster OEP5 and blue arrows highlight overlapping LHS cluster between EPP and Estrogen (OEP6). (D) Bar plot showing percentage of cells per condition in each cellular cluster. (E) Module score analysis for markers from cluster OE6 in OP clusters. The blue arrows point to clusters with a high score for OE6 markers, clusters OP2, OP4, OP7, OP8, OP9 and OP10. (F) OIP clusters split by condition, highlighting a cellular state enriched in EPP-treated organoids (blue arrow). (G) Regulon analysis of clusters expanded in EPP.

For data presented on Figure 4, scRNAseq profiles of untreated human organoids, and pregnancy hormone treated ones (10 days and 21 days of EPP treatment), low quality cells were removed, yielding a total of 14,621 cells from organoids without treatment, 5,888 cells from organoids at 10 days of EPP treatment, and 8,167 cells from organoids at 21 days of EPP treatment, respectively, which were utilized on further analysis.

**Figure 4.**
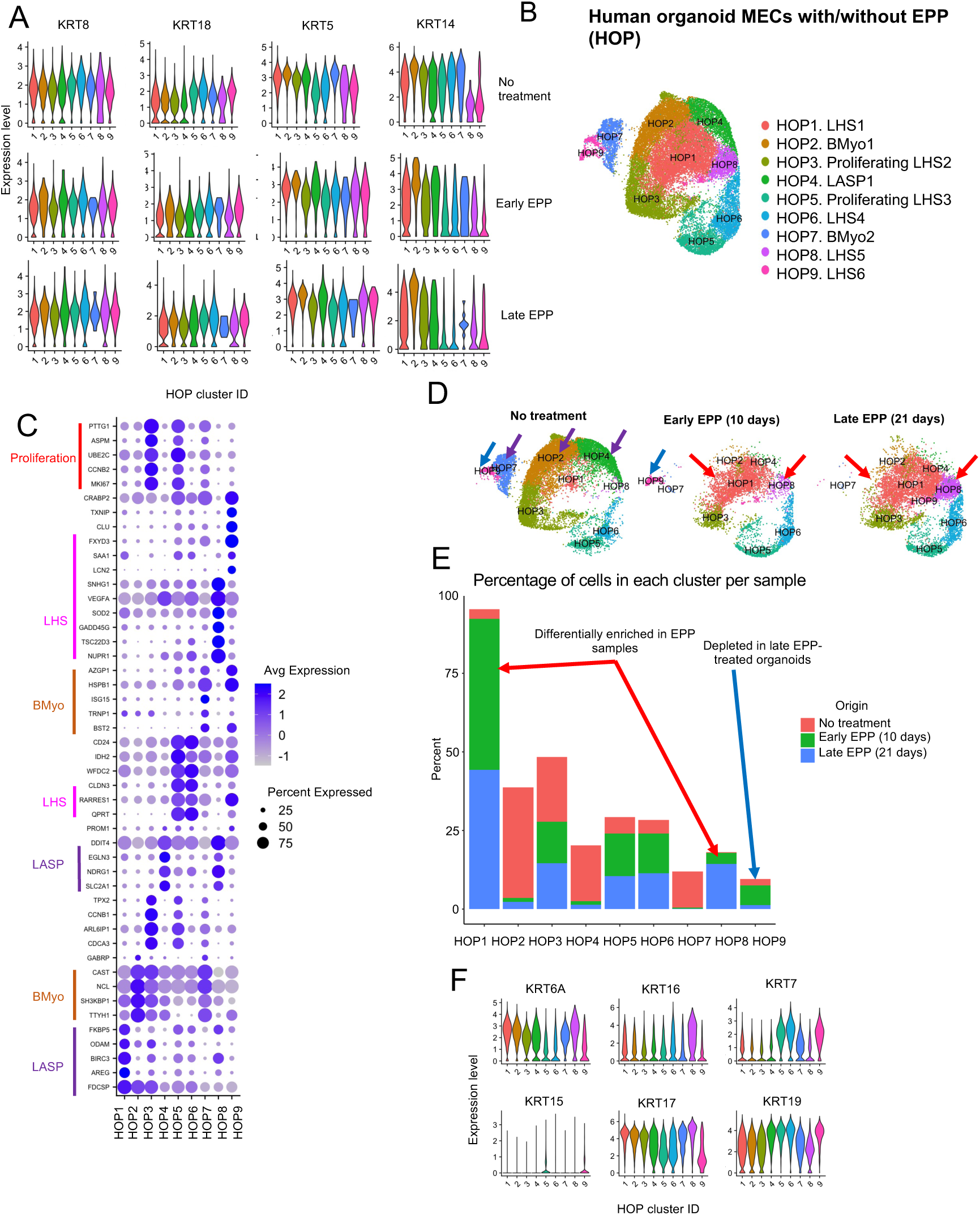
Analysis of scRNA-seq data from human organoid MECs treated with EPP. (A) Violin Plots showing the expression of cytokeratins used to classify luminal and basal populations within each HOP cluster, divided by condition (no EPP treatment, early EPP treatment and late EPP treatment. (B) Resulting clusters for human organoid MECs with and without EPP treatment (HOP), along with cluster identities based on expression of previously described markers from intact human MECs and top differentially expressed genes (DEGs) per cluster. (C) Dotplot for top DEGs per HOP cluster. Clusters are organized based on dendrogram relationships. (D) HOP clusters split by condition. The purple arrows highlight clusters enriched in organoids without treatment, and red arrows highlight clusters enriched with EPP, independent of the amount of time with EPP treatment. (E) Bar plot showing percentage of cells per condition in each HOP cluster. The red arrow highlights HOP2, most notably enriched in EPP treatment at 10 and 21 days. The blue arrow highlights HOP9, which is depleted during late (21 days) EPP treatment. (F) Violin Plots showing the expression of additional cytokeratins used to classify luminal and basal populations within each HOP cluster.

For data investigating similarities across species, murine genes were converted into their human orthologs before scRNAseq data integration (Zilionis et al., 2019). This approach yielded a total of 7 clusters for our untreated human and murine organoid (UHM) comparison, with 14,621 cells from humans and 10,508 cells from murine organoids. Similarly, for our murine and human organoids with pregnancy hormones (PMH) comparison, the aforementioned approach yielded 5 clusters, with 14,055 cells from humans and 16,463 cells from murine MEC organoids.

Pathway analysis was performed using Gene Set Enrichment Analysis (GSEA) v3.0 and with the Molecular Signatures Database (MSigDB) Hallmark Terms (Liberzon et al., 2015; Mootha et al., 2003; Subramanian et al., 2005). This database was selected with the purpose of obtaining an overview of the processes each cellular cluster was undergoing. The resulting hallmark terms were further filtered based on their nominal (nom) p-value (<0.05), with the purpose of only showing significant terms per cluster and/or condition. The -log(nom p-value) for each hallmark term was calculated so that these could be visualized based on significance. On Figure 1, given that most clusters had similar signature gene modules, differentially expressed pathways with an adjusted p-value of <0.06 were kept for further analysis.

## Results

### 1. Determining the cellular landscape of murine mammary organoids

Previous studies have demonstrated the capacity of human MEC-derived organoids in retaining *in vivo* lineages (Gray et al., 2022; Rosenbluth et al., 2020). Further studies have demonstrated the feasibility of murine organoid systems to recapitulate parity-associated phenotype, such as expressing milk associated proteins and parity-associated epigenomic signatures (Ciccone et al., 2020; Sumbal et al., 2020). However, the heterogeneity of murine organoids cultures, and how it recapitulates the heterogeneity of intact tissue remains to be elucidated. In order to assess the cellular molecular heterogeneity of mammary organoids, we sequenced mammary organoids cells, derived from nulliparous female mice using the 10X Chromium platform.

Utilization of previously defined markers for lineage commitment in intact mammary tissue (Henry et al., 2021) allowed for robust classification of mammary organoid (MO) cell types. Such analysis identified 5 populations of luminal epithelial cells, marked by the expression of both cytokeratin 8 and 18 (*Krt8*/*Krt18*) markers, (MO1, MO2, MO3, MO5 and MO7), one population of basal myoepithelial (BMyo) cells defined by the expression of cytokeratin 5 and 14 (*Krt5*/*Krt14*) markers (MO4), and a cluster of cells expressing both luminal and myoepithelial markers (MO6) (**Fig. 1A** and **S1A**).

Further gene expression analysis shed light into the lineage subtypes of each cellular cluster. Expression of hormone receptors such as progesterone Receptor (*Pgr*), prolactin Receptor (*Prlr*) and estrogen Receptor a (*Esr1*) defined luminal populations of hormone sensing (LHS) cells MO2 and MO7 (**Fig. 1A** **and S1A**). Cluster MO1, MO3 and MO5 were defined to have a luminal adaptive secretory precursor fate (LASPs), given higher expression levels of genes linked to milk synthesis, such as Casein 3 (*Csn3*), and Lactalbumin Alpha (*Lalba*) (Bach et al., 2017; Saeki et al., 2021) (**Fig. 1A** **and S1A**). Luminal cluster MO5 (LASP) and MO7 (LHS) were also characterized by the expression of genes associated with highly proliferative gene signature such as Marker of Proliferation Ki-67 (*Mki67*), Ubiquitin Conjugating Enzyme E2 C (*Ube2c*), DNA Topoisomerase II Alpha (*Top2a*), thus defined as proliferative cellular states (**Fig. S1A**). Further cell cycle scoring analysis confirmed that epithelial cells from both MO5 and MO7 clusters were predominantly at G2M and S-phase stages of cell cycle, thus supporting that several luminal subtypes assume a proliferative state in organoid cultures (**Fig. S1B**).

Our analysis also identified molecular states of less differentiated cell types. We found that cells from clusters MO1, MO3 and MO5 were characterized by expression of luminal progenitor genes FXYD domain-containing ion transport regulator 3 (*Fxyd3*), Cluster of differentiation 14 (*Cd14*) and Claudin-3 (*Cldn3*) (Asselin-Labat et al., 2011; Coradini et al., 2014; Shehata et al., 2012; H. Wang et al., 2019) (**Fig. S1A**). Cluster MO3 cells also expressed genes associated with milk synthesis WAP four-disulfide core domain protein 18 (*Wfdc18*) and Mucin-15 (*Muc15*), suggesting a secretory progenitor state for cells from this cluster (Pal et al., 2021; C. Shao et al., 2021) (**Fig. S1A**). Our analysis also supported a mixed lineage state to cells from cluster MO6, given the expression of markers for both luminal and basal lineages, as well as Galectin-1 (*Lgals1*) expression, a previously identified marker for mammary stem cells (Soady et al., 2015). To further support the identification of MO6, we obtained a list of genes associated with the term “mammary stem cell” from Enrichr (Chen et al., 2013), and then assigned a score to each cell within MO clusters for the expression of these stem-associated genes (**Table S1**). We observed that most MO clusters expressed the aforementioned genes, including MO6, further suggesting a stem-like state for most MECs in culture, and a mixed lineage phenotype present amongst these progenitors (**Fig. S1C**). Moreover, the suggested lineage identities of all organoid epithelial cell types were supported by the utilization of ternary plot analysis, which indicated a less committed lineage distribution of progenitor (MO1, MO3, MO5) and mixed lineage clusters (MO6), while more differentiated luminal (MO2 and MO7) and BMyo (MO4) clusters aligned alongside their predicted lineage identities (**Fig. 1B**).

We next investigated which molecular signatures were enriched in each cluster identified in organoid cultures. While clusters MO5 and MO7 were enriched with pathways associated with cell division, cells from cluster MO2 were marked by processes associated with LHS cells, thus collectively supporting their above assigned cellular states (**Fig. 1A-B, S1A and S1D-E**). Accordingly, the BMyo state of cells from cluster MO4 were further supported by the enrichment of genes associated with myogenesis and EMT-like processes (Ingthorsson et al., 2015). Cells from LASP cluster MO1 were significantly enriched for terms involved in hypoxia. However, when considering hypoxic genes detected in our dataset, we found that most of these were involved in milk-synthesis, such as *Lalba* and *Aldoc*, supporting a secretory classification (**Table S2**) (Bach et al., 2017; Rudolph et al., 2007; Saeki et al., 2021). Mixed lineage cells in MO6 were enriched for terms similar to BMyo cluster MO4, as well as expressing genes involved in p53 signaling and coagulation. Adequate p53 signaling has been implicated in mammary tissue homeostasis during development (Dusek et al., 2012). Moreover, genes involved in coagulation were also implicated in EMT processes, such as Fibronectin 1 (*Fbn1*) and Kallikrein-related peptidase 8 (*Klk8*) (**Table S2**) (Bahcecioglu et al., 2021; Hua et al., 2021). Interestingly, cells from cluster MO3, classified as LASPs, did not show enrichment for specific terms in relation to all other cell types, thus suggesting an organoid cellular state that shares transcriptional signatures with all other cellular clusters. We compared MO3 to MO1, in order to explore how early LASPs differ from those that are *Lalba*+ (**Fig. S1E**). The aforementioned analysis revealed that cells in MO3 are enriched for genes associated with apoptosis and EMT, both which have been associated with undifferentiated process in mammary epithelial cells and thus suggest an increased plastic state for cells in MO3 compared to cells in MO1 (C.-W. Li et al., 2012).

It is possible that a less defined cellular identity of organoid cells could represent changes induced by *ex vivo* culturing that alters molecular signatures that define MECs cell types. In fact, transcriptional and cellular profiles of human breast organoid cultures from normal and cancer derived tissue suggest that over-time, culturing conditions can induce gene expression and lineage marker changes (Bhatia et al., 2022; Gray et al., 2022). In order to define the culture-induced changes to mammary organoid cultures, we integrated previously published scRNA-seq datasets to map epithelial cells from nulliparous mice mammary tissue to our analysis, to define the similarities and differences across *in vivo* and *ex vivo* MECs (Bach et al., 2017; Henry et al., 2021). This analysis yielded 6 epithelial clusters, including those of luminal fate (OIM1, OIM2, OIM3, OIM5 and OIM6) or BMyo lineage (OIM4) (**Fig. 1C** **and S1F**). Overall, the majority of clusters defined on organoid cultures were also represented in intact mammary tissue, with the exception of cluster OIM6, which was markedly expanded in libraries prepared from organoid conditions (**Fig. 1D-E**). Interestingly, global expression hierarchical relationship across all clusters (dendrogram), indicated a closer relationship between cluster OIM1 (LHS cells) and OIM6, which lacks the expression signature of hormone-responsive cells (**Fig. S1F**). Conversely, OIM6 expressed elevated levels of genes in LASP cellular states such as *Csn3*, *Trf*, and *Gm42418*, in comparison to cells from OIM1 cluster, suggesting an expression signature of a not fully defined luminal state (**Fig. S1F**). In fact, our analysis indicated that OIM6 cells are positioned in an intermediary state, right in between LHS cluster (OIM1), and LASP clusters (*Lalba*+ OIM2, and *Aldh1a3*+ OIM3), further suggesting a transitional LASP state (**Fig. 1F**). GSEA for hallmark terms revealed that organoid-exclusive cluster OIM6 was significantly enriched for terms involving apoptosis and EMT, similar to what we observed in cells within MO3, thus further suggesting the presence of organoid cells with early progenitor phenotypes in culture (**Fig. 1G** **and S1D**) (C.-W. Li et al., 2012).

Overall, our initial mapping of molecular and cellular makeup of mammary-derived organoid cultures illustrates aspects of *ex vivo* models that resemble intact mammary tissue, while highlighting those that are induced by several of the stimuli of a culturing system.

### 2. Characterizing the effects of estrogen treatment on mammary-derived organoid cultures

Puberty represents the first key signal post-birth that drives mammary tissue expansion and MEC lineage differentiation, with increased levels of estrogen regulating cell-to-cell signaling, immune modulation, and transcription regulation (Rusidzé et al., 2021; Tower et al., 2022; Vasquez, 2018). Once developed, physiological levels of estrogen sustain mammary tissue homeostasis, with cyclical cellular dynamics throughout the estrous cycle further influencing MEC differentiation and proliferation (Pal et al., 2017). Yet, the necessity and effects of estrogen supplementation for the growth of mammary organoid cultures has not been fully characterized.

With the purpose of determining the effects of estrogen on gene expression, growth, and cellular heterogeneity, we set out to characterize mammary organoids treated with two concentrations of 17-β-Estradiol, 66.6 ng/mL (i.e. “high estrogen”) and 33.3 ng/mL (i.e. “low estrogen”) (referred hereafter as OE) (**Fig. S2A**). The higher estrogen concentration aimed to replicate levels found during peak estrogen production, such as during pregnancy, while the lower concentration sought to mimic physiological levels of the hormone. Our analysis identified several clusters in all conditions, spanning BMyo fates (OE4), LHS states (OE6 and OE9), LASP subtypes (OE1, OE2, OE3, OE8), and cells expressing both luminal and basal cells lineage markers, referred hereafter as mixed lineages subtypes (OE7) (**Fig. 2A-B**). We also identified cellular clusters marked by the expression of proliferation markers, encompassing LHS (OE9), and LASP (OE5) luminal states (**Fig. 2A** **and S2B-C**).

Further analysis of cell population distribution across organoid conditions indicated a few cellular clusters biased to specific datasets (**Fig. 2A**).We found a subtle decrease on the abundance of LASPs (cluster OE3) in organoid conditions supplemented with estrogen, perhaps suggesting that luminal progenitor differentiation in response to increased levels of estrogen can also be observed in organoid cultures (Basak et al., 2015) (**Fig. 2A**). Depletion of mixed lineage cells (cluster OE7) was also observed in organoid cultures treated with estrogen, supporting the suggestion that estrogen supplementation may be inducing the differentiation of immature cell types, as is observed *in vivo* (Simões & Vivanco, 2011). Interestingly, none of these cell types express hormone genes, thus suggesting a possible indirect effect of estrogen on their homeostasis/differentiation (**Fig. S2B**) (Sleeman et al., 2006). We also identified alteration to cluster of LHS cells (OE6), thus validating that expression of hormone responsive genes in subtypes of organoid cells are linked with cellular expansion in response to increased estrogen levels (**Fig. 2A-B, and Fig. S2B**) (Feng et al., 2007).

Given that our observations indicated that estrogen supplementation of organoid cultures can influence cellular populations independently of the expression of hormone-associated genes, we next decided to investigate global gene expression alterations across all organoid clusters. For this, we started looking at gene expression alterations across organoids without hormone treatment and those treated with low levels of estrogen, given that all identified clusters are represented in both conditions (**Fig. 2C**). Our analysis identified that clusters defined to have an LHS identity where the ones with the most alterations to enriched pathways in response to estrogen treatment, with proliferative LHS cells (OE9) demonstrating selective enrichment for processes associated with estrogen response (early and late) and K-ras signaling, a pathway previously associated with estrogen receptor signaling (Dischinger et al., 2018) (**Fig. 2C**). Conversely, the population of LHS cells expanded in response to estrogen levels (OE6) was selectively enriched for pathways associated with reactive oxygen response and genes that downregulate UV responses, both potential antioxidant pathways also described to be regulated by estrogen (Caldon, 2014; Halliday, 2010). This observation was in marked contrast to the observed in subtypes of LASP cells (OE1), which increased expression of UV responses was linked with estrogen treatment. Both OE6 and OE9 clusters were also enriched for pathways related to hypoxia, thus suggesting the diverse gene regulation modules regulated by estrogen on LHS cells (**Fig. 2C**).

In addition to pathways that were shared with cell types defined as LHS, estrogen treatment of organoids induced enrichment to specific pathways in a hormone expression independent manner. For example, enrichment for TNF-⍺ signaling via NF-κB pathways was observed in both LASPs (OE1, OE5) and LHS cells (OE6, OE9). In addition, mixed lineage cells (cluster OE7) and those of proliferating LHS cells (OE9) were exclusively enriched with genes associated with myogenesis, a process that can either be suppressed or activated by estrogen levels on cellular contact dependent fashion, thus suggesting a deeper level of estrogen related signals, that can be recapitulated in *ex vivo* organoid cultures (**Fig. 2C**) (Mallepell et al., 2006; Ogawa et al., 2011; Strum, 1978). Another controversial signal pathway regulated by estrogen, EMT, was also observed on luminal fate clusters mostly affected by estrogen supplementation, including LASPs (cluster OE3), LHS cells (OE6), and proliferative LHS cells (OE9), an observation that may link EMT with loss of cell plasticity, and estrogen-induced differentiation (**Fig. 2C**) (Guttilla et al., 2012; Wahl & Spike, 2017). In fact, the only statistical significantly enriched pathway downregulated by estrogen was associated with cMYC regulated processes in transitional LASPs (cluster OE3), a signal that is essential to keep immature properties of mammary epithelial cells (**Fig. 2C**) (Poli et al., 2018). Despite the findings described above, low levels of estrogen did not result in the significant enrichment of pathways in clusters of cells with BMyo fate (OE4), or certain LASP populations (OE2 and OE8), suggesting that subtypes of MECs that lack the expression of hormone genes are less affected by female hormones. Nonetheless, high levels of estrogen did enrich the aforementioned non-hormone sensing subtypes for oxidative phosphorylation-associated genes, indicating that non-hormone sensing cells are still capable of responding to hormones, at a lesser degree (**Fig. 2D**).

To assess how the regulatory networks modulating processes in each cellular sub-type might be affected by estrogen, we calculated the regulons with the highest specificity scores (RSS) for each of the OE clusters and segregated them by condition (i.e., untreated, low estrogen treatment and High estrogen treatment) (**Fig. 2E**). Our analysis identified a series of regulons that defined overall cellular states. Luminal epithelial cells with higher proliferative states (OE5 and OE9) were defined by programs regulated by general cell cycle regulators including Rad21, Ybx1 and Chd2, suggesting a proliferation regulatory network independently of the lineage state (Jurchott et al., 2003; Mills, 2017; Sun et al., 2023). Clusters of luminal and myoepithelial cellular states were enriched for several known lineage defining regulatory networks, with LHS clusters (OE3, OE6, and OE9) sharing programs regulated by Gata3, Jun, Foxa1, Tead2, and Jund, while BMyo-biased cell clusters (OE4 and OE7) were marked by basal-biased transcription regulators such as Runx2, Trp63, Tead1, and Sox4 (Assefnia et al., 2014; Kendrick et al., 2008; Knight et al., 2008; McDonald et al., 2014; Roukens et al., 2021).

Several of the proposed luminal progenitor cellular states (OE1, OE2 and OE8) shared several transcription modules, including those regulated by Nf1, Cebpb, Sox10, Sox5, Trsp1, all factors previously defined to regulate progenitor cell activity during mammary gland development (Cornelissen et al., 2020; Dischinger et al., 2018; LaMarca et al., 2010; Y. Liu & Guo, 2021; Mertelmeyer et al., 2020) (**Fig. 2E**). Interestingly, BMyo cell cluster OE4 and mixed lineage cell cluster OE7 shared similar transcription networks, with the exception of programs regulated by Creb3, which was also enriched in clusters defined to have LASP signatures (OE2) and LHS state (OE3) (**Fig. 2E**). In fact, Creb3 has been suggested to have increased activity in cells undergoing luminal-basal cellular plasticity in response to high levels of Sox9, thus supporting the suggested mixed lineage state of cells from cluster OE7 (Christin et al., 2020). These observations suggest that mixed lineage cell types have a transcriptional identity that resembled basal states closely, with discrete alterations to luminal-biases programs.

We also identified estrogen-induced changes to transcriptional programs, encompassing both alterations to several lineage restricted programs, and those spanning several cellular states. While estrogen treatment induced an overall decrease of lineage transcriptional programs signal in LHS cluster OE3, analysis of BMyo cells (cluster OE4) demonstrated a bimodal change of basal transcription programs, with the enrichment of luminal-basal plasticity regulators such as Creb3, Tfe3, and Sox4, and partial loss of programs controlled by Relb, which has been reported to repress the expression of estrogen Receptor α (ERα), and Zfp358, which has been noted to decrease with Estradiol treatments (Vydra et al., 2019; X. Wang et al., 2009) (**Fig. 2E**). Further regulon analysis of LASP cell types indicated a further enrichment of lineage specific transcriptional programs in specific cellular clusters (OE2, and OE5) (**Fig. 2E**). Collectively, these analyses suggest that estrogen treatment impacts the lineage programs of specific luminal and myoepithelial cell types, thus indicating cell types that are the most responsive to increased levels of female hormone.

Interestingly, our investigation identified a group of transcription programs that were altered in a lineage-independent fashion in response to increased estrogen levels, which included programs regulated by Edr, Ehf, Creb5, Tfdp2, Elf2, Sin3a, E2f4, previously linked with regulating the cell cycle, especially in promoting cell growth and proliferation (Hadsell et al., 2015; Johnson et al., 2016; W. Li et al., n.d.; Luo et al., 2020; Qiu, Morii, et al., 2008). Interestingly, Ehf has been reported to increase when mammary stem cells begin the process of differentiation, suggesting a role for estrogen in the maturation of hormone receptor negative MECs (Williams et al., 2009). Our analysis indicated gains and losses of these regulon activities across all identified cellular states, thus further illustrating the complex effect of estrogen on regulatory process of all subtypes of mammary epithelial cells (**Fig. 2E**).

### 3. Pregnancy hormones exposure, cellular states, and gene expression

Mammary organoid systems have been previously optimized to mimic aspects of pregnancy-induced development of the gland, such as branching and production of milk-associated proteins, involution-like processes, and mechano-regulated actions of lactation (Ciccone et al., 2020; Stewart et al., 2021; Sumbal et al., 2020). Yet, it is unclear whether mimicking pregnancy-induced changes *ex vivo* drives cellular and transcription alterations such as those that take place *in vivo*. Therefore, we set out to characterize mammary organoid cultures, grown with a combination of estrogen, progesterone, and prolactin (EPP) hormones (referred hereafter as OP) using scRNA-seq approaches. Our analysis identified clusters present in both untreated and EPP-supplemented conditions, encompassing cellular states of LASP fate (OP1, OP3, and OP6), BMyo lineage (OP5), in addition to lineages more abundant in untreated organoids (LHS clusters OP4, OP9, and mixed lineage cluster OP7), and those expanded in EPP-treated conditions (LHS clusters OP2, OP8, and mixed lineage cluster OP10) (**Fig. 3A-B** **and Fig. S3A**). Amongst these clusters, we identified highly proliferative cells in both conditions (OP6) (**Fig. 3A-B** **and Fig. S3B-C**).

We next defined the pathways differentially expressed in each of the identified cellular clusters in response to treatment with EPP. Across the cellular clusters that were present in both untreated and EPP-treated conditions, which encompassed hormone negative cell types (OP1, OP3, OP5, and OP6), we found clusters with no statistically significant enrichment for specific terms (OP1, and OP3, LASP identity), indicating cellular stages that were minimally affected by pregnancy hormones (**Fig. S3D**). Conversely, clusters identified as BMyo lineage (OP5) and proliferating LASPs (OP6) were enriched for term that were related to their lineage specific developmental state (such as myogenesis and EMT for OP5) (Mallepell et al., 2006; Ogawa et al., 2011; Strum, 1978), or cellular state (mitotic spindle and G2M checkpoint for OP6), suggesting that similarly like estrogen alone, pregnancy hormones can induce indirect transcription changes in hormone negative cells (**Fig. S3D**).

Interestingly, clusters biased towards samples without hormone treatment (OP4, OP7, OP9) and those more abundant in EPP-treated samples (OP2, OP8, OP10), represented very similar cellular identities, with LHS cells (OP4, OP9, OP2, OP8) and mixed lineage fate (OP7, OP10), suggesting that pregnancy hormones act on cellular states fully present in conditions without hormone treatment (**Fig. S3A-B and Fig. S3B**). In fact, LHS clusters OP4 (untreated condition), and OP2 (EPP condition) where enrichment for similar pathways such as estrogen response and hypoxia, with the exception of untreated cluster OP4, that was also enriched for pathways regulated by p53 (**Fig. S3D**). Interestingly p53 pathways have been associated with acquisition of a senescent-like state, which can be modulated by pregnancy hormones (Feigman et al., 2020). Additional LHS clusters OP9 (untreated condition) and OP8 (EPP condition) were enriched with terms associated with cell division, thus validating our initial classification of these clusters as proliferative states (**Fig. S3C-D**).

Moreover, pathways associated with myogenesis and EMT were both enriched in mixed lineage clusters OP7 (untreated condition) and OP10 (EPP condition), with the specific enrichment of p53 pathways in cells from untreated conditions, supporting the suggestion that pregnancy hormones may suppress similar pathway in different cellular states. The hormone expression on cells from OP10 cluster was linked to an enrichment of estrogen response and hypoxia, a pathway also associated with pregnancy signals, thus suggesting that hormone regulated pathways are also synchronized in more immature cell types (**Fig. S3D**) (Y. Shao & Zhao, 2014). Collectively, this pathway analysis mapped the transcriptional alteration to organoid cultures in response to pregnancy hormones.

Given that our previous analysis demonstrated that supplementation of estrogen induced molecular and cellular change on organoid cultures, we next investigated whether the pregnancy-hormones induced changes were driven collectively by estrogen, progesterone and prolactin, or rather represent alterations regulated by estrogen alone. In doing so, we compared the transcription and cellular dynamics of estrogen treated organoids (OE), with those present in organoids cultured with EPP (OP). This analysis yielded 9 clusters (referred thereafter as OEP), encompassing populations of BMyo cells (OEP4), progenitor and mixed lineage cells (OEP2, OEP6, OEP8, OEP9), LHS (OEP3, OEP5, OEP7), and LASP cells (OEP1) (**Fig. 3C-D** **and S3E**). We found that that the majority of clusters are present in both culturing conditions, with the exception of 2 populations of LHS cells (clusters OEP3 and OEP7) which were exclusive to conditions treated with EPP, thus indicating cellular dynamics that only take place when estrogen, progesterone and prolactin are in place (**Fig. 3C-D** **and S3E**).

To more accurately identify estrogen alone induced changes in organoid cultures grown with EPP, we defined gene core signatures that defined the previously identified estrogen-induced cluster OE6 of LHS cells (**Fig. 2**), and asked whether these markers were also upregulated in population of cells grown with EPP (**Table S3,** **Fig. 3E**). We observed that a large quantity of LHS cells in clusters enriched with EPP (OP2 and OP8) appeared to score highly for OE6 markers compared to other clusters, with only a small portion of LHS cells in clusters enriched without treatment also scoring highly for OE6 markers (OP4 and OP9) (**Fig. 3E**). These results indicate that estrogen alone induces an expansion of LHS cells, while progesterone and prolactin enhances transcriptomic differences between LHS cells without hormone treatment and those treated with EPP. We also observed that more mixed lineage cells in cluster OP10 enriched with EPP scored highly for OE6 clusters compared to cluster OP7 of mixed lineage cells enriched with no hormone treatment. Therefore, given that only a portion of mixed lineage cells in OP7 score highly for OE6 markers and are depleted with EPP, this suggests that these OP7 cells are estrogen responsive and obtain an EPP-exclusive signature upon exposure to EPP (**Fig. 3A** **and** **Fig. 3E**).

Our cellular and differential gene expression analysis identified many pathways that are activated specifically in response to organoid cultures treated EPP (**Fig. S3D**). Our initial analysis indicated that untreated organoids bear similar cell types and transcriptional output than MECs directly extracted from mammary tissue (**Fig. 1D**). Therefore, in order to assess whether clusters identified in our hormone treated culturing system were also represented during pregnancy in mice, we performed a data integration analysis, comparing organoid datasets with publicly available profiles generated from MECs during distinct stages of pregnancy (gestation, lactation, and involution) (Bach et al., 2017).

With this approach, we found several clusters with similar representation in all datasets, including those made up of LASPs (OIP1), two populations of BMyo cells with slightly less abundance in samples from mammary tissue during involution (OIP3 and OIP4), and those with more abundance during lactation (OIP4) (**Fig. 3F** **and Fig. S3F-H**). Cluster OIP6, identified as a population of LHS cells, was also detected across all datasets, but with overall low abundance (**Fig. 3F** **and S3H**). Collectively, this initial analysis identified an array of cellular states, that are present in the mammary tissue across the pregnancy cycle, and sustained in organoid cultures grown with EPP.

This analysis also identified populations of cells biased towards specific stages of pregnancy-induced development, and that were also present in organoid cultures treated with EPP. EPP-treated cultures were enriched with specific populations of LHS cells (OIP2), which were also present in mammary tissue during involution, perhaps indicating cellular populations that are expanded in response to pregnancy hormones, and remain present even when signals of pregnancy are gone (**Fig. 3F** **and S3H**). We also identified a population of proliferating LASPs (cluster OIP5), which were more abundant in EPP-treated organoid cultures and in mammary tissue during gestation, suggesting populations of cells that are activated by hormones early during the pregnancy cycle (**Fig. 3F** **and S3G-H**).

Interestingly, our analysis also identified proliferating LASP states in organoids without hormone treatment (MO4), and as well in those treated with estrogen levels (OE5), and EPP (OP6), suggesting that organoid culturing conditions may in general provide the signals for active proliferation of cell types (**Fig. S1B**, **S2C** and **S3C**). In fact, the cluster with greater bias towards EPP-treated organoids, in comparison to mammary tissue during pregnancy cycle, represents a population of proliferative LHS cells (OIP7), detected in untreated and hormone treated organoid conditions, thus further suggesting that such culturing conditions support proliferative states independently of hormone supplementation (**Fig. 3F** **and S3I**). However, specific populations of proliferative LHS cells were detected in response to EPP treatment (OP9), thus suggesting an additional level of cell proliferation activation in response to specific stimuli (**Fig. S3C**). Collectively, this organoid versus normal tissue comparative analysis further illustrates similarities across the systems, therefore lending significance to the utilization of organoids to understand pregnancy-induced mammary development.

It is very well established that pregnancy hormones, such as estrogen, progesterone and prolactin, can act directly on chromatin, thus controlling gene expression, in addition to modulate additional downstream signals of gene regulation. Therefore, we next set out to define the pool of enrichment for regulons across all clusters, in response to pregnancy hormones (**Fig. 3G**). This approach allowed the identification of transcription programs of LHS cells, including core programs enriched in all LHS cells types (OP4, OP9, OP2, OP8), such as Runx1, Tbx3, Foxa1, in addition the enrichment of Gata3, Junb, Irf3 programs in cellular states more abundant in untreated conditions (OP4 and OP9), and programs regulated by Stat5a and Stat5b, Pgr, Jun, Klf6, Tfcp2l1, Myb, Spdef which were enriched in EPP-treated conditions (Arendt & Kuperwasser, 2015; Balogh et al., 2006; Bernardo et al., 2010; Bjornstrom et al., 2001; Cicatiello et al., 2010; Davenport et al., 2003; de Assis et al., 2013; Eeckhoute et al., 2007; X. Liu et al., 1997; Otto et al., 2013; Quintana et al., 2011; van Bragt et al., 2014; Ye et al., 2020) (**Fig. 3G**).

In addition, we identified regulons that defined proliferative cellular states (clusters OP2, OP8, and OP10), including programs controlled by Yy1 and Gtf2f1, which were also enriched in non-proliferative clusters OP4, OP7, and OP2, specifically in EPP-treated conditions, suggesting a discrete increase on cell division in response to pregnancy hormones (Rizkallah & Hurt, 2009; Zhang et al., 2022, p. 2) (**Fig. 3G**). Regulons for both BMyo cells (clusters OP5, OP7, OP10) and luminal progenitor cells (clusters OP1, OP3) were defined by classic lineage defining clusters, with the enrichment of Trp63, Tead1, Relb, and Sox4 in clusters OP5, OP7, OP10, and Trsp1, Cebpb, Sox10, Egr1 controlled programs enriched on clusters OP1, OP3 (Assefnia et al., 2014; A. E. Casey et al., 2018; Cornelissen et al., 2020; Dravis et al., 2015; Gu et al., 2021; Knight et al., 2008; Roukens et al., 2021; X. Wang et al., 2009) (**Fig. 3G**). Our analysis also identified several regulons enriched in EPP-treated condition, in a non-lineage fashion, which included programs regulated by Tcf7l2, Phfl8, Sp1, Arntl, Nfkb1 in LHS cells (cluster OP4), BMyo subtypes (clusters OP5 and OP7), and in LASP types (clusters OP1, OP3 and OP6), suggesting mechanisms that regulate pregnancy-induced responses in all major mammary epithelial cell types (T. M. Casey et al., 2014; Cho et al., 2020; Dong et al., 2016; Gómez-Chávez et al., 2021; Schanton et al., 2017) (**Fig. 3G**).

Overall, our approach to profile of molecular mechanisms regulating cellular states and pregnancy-induced development of organoid systems provide a solid framework for the identification of novel master regulators of mammary lineage identity and development.

### 4. Defining the molecular alterations induced by pregnancy hormones on human MEC-derived organoids

The current understanding of tissue alterations in response to pregnancy signals is largely biased towards the investigation of molecular and cellular dynamics in rodent models. Our above-mentioned findings suggest that the utilization of organoid cultures represent a suitable system to model, at least partially, the response of MECs to female puberty and pregnancy hormones. Given that normal, human breast tissue has been utilized for the development of organoid systems (Bhatia et al., 2022; Gray et al., 2022; Rosenbluth et al., 2020; Sachs et al., 2018), we next decided to test their response to supplementation with EPP.

In doing so, we utilized an already established and characterized normal breast organoid culture, generated from breast specimens from women undergoing cosmetic reduction mammoplasty (Bhatia et al., 2022). Human organoid cultures were treated with the same concentration of estrogen, progesterone and prolactin that was employed for mouse mammary organoids, given that human MECs have been shown to engage on pregnancy-induced development in response to pregnancy in mice (Kuperwasser et al., 2004). Pregnancy-induced development was confirmed with the quantification of CSN2 mRNA levels, previously described to increase in response to pregnancy hormones (Maningat et al., 2009; Rijnkels et al., 2013). Our qPCR analysis indicated significant increased levels of pregnancy-specific *CSN2* mRNA in contrast to *CSN3* levels, starting on day 10 after EPP treatment, a response that was sustained up to 21 days of culturing (**Fig. S4A**). This observation was confirmed by the detection of milk-associated proteins in human organoid cultures treated with EPP for ∼21-25 days, thus further supporting that such approach promotes pregnancy-associated changes (**Fig. S4B**). Therefore, we utilized the same culturing conditions for the generation of scRNAseq profiles of untreated and pregnancy-hormone treated human mammary organoids.

Characterization of lineage identity, utilizing classic markers of luminal and basal breast epithelial cells, indicated that the majority of cells in untreated and EPP-treated organoid cultures bear both luminal and basal traits, defined by the expression of *KRT8*, *KRT18*, *KRT5* and *KRT14*, suggesting that independent of treatment, established human breast organoid system have a more generalized mix-lineage signature (**Fig. 4A**). This observation agrees with previous studies describing that human breast organoid systems assume a more basal-like cellular phenotype after several culture passages, with consecutive loss of hormone receptor expression (Bhatia et al., 2022). Interestingly, analysis of *KRT14* mRNA levels indicated clusters with high, low, and moderated levels of expression, suggesting that at least 3 epithelial lineages could be delineated (**Fig. 4A**). Therefore, and with the goal to define cellular states of established human breast organoid cultures, we employed an approach that utilized top differentially expressed genes across all clusters, and markers previously utilized to define human MECs identities (Henry et al., 2021) (**Fig. 4B-C**).

With this approach, we identified a unique population of LHS progenitor cells (HOP3), present in all culturing conditions, which expressed low levels of *KRT14*, and were defined by the expression of progenitor markers *FDCSP* and *ODAM* (Kestler et al., 2011; McMullen & Soto, 2022), pregnancy hormone regulated genes (*BIRC3*) (LaMarca & Rosen, 2007), and proliferating cell markers such as *MKI67, CCBN2* and *PTTG1* (Neubauer et al., 2011; Wei et al., 2013) (**Fig. 4C-E** **and S4C**). Further analysis, identified cell populations that were biased towards untreated human organoid samples, encompassing 2 populations of BMyo cell types (HOP2 and HOP7) which expressed high levels of *KRT14* mRNA, and high levels of basal-like cell identity such as *TTYH1, BT2, ISG15,* and *SH3KBP1* (Bolado-Carrancio et al., 2021; Klinke & Torang, 2019; Sayeed et al., 2013). Interestingly, the expression of many of these genes were elevated across additional clusters, further supporting a more basal-like phenotype to human breast organoid cultures (**Fig. 4C**). We also identified a population of LASP cells (HOP4) to be more abundant in organoids without hormone treatment, and marked by the expression of lactogenic-associated genes such as *SLC2A1*, *NDRG1* and *EGLN3* (Pal et al., 2013; Williams et al., 2009; Zhao & Keating, 2007) (**Fig. 4C-E**). Collectively, our analysis suggests the existence of population of cells that are present in human breast organoid conditions, and that are negatively impacted by the presence of pregnancy hormones.

We next focused on the characterization of cellular clusters biased to EPP-treated conditions. This approached identified cell types spanning a series of LHS states, mostly marked by lower levels of *KRT14* mRNA, and variable levels hormone responsive genes such as *BIRC3, RARRES1,* and *NUPR1* (HOP1, HOP3, HOP5, HOP6, HOP8 and HOP9) (Bhat-Nakshatri et al., 2021; Hollmén et al., 2015; Neubauer et al., 2011; Zhou et al., 2014) (**Fig. 4C-E**). Here we defined populations of LASP cells expressing estrogen/progesterone-associated genes such as *AREG, ODAM,* and *FKBP5* (Cai et al., 2020; Habara et al., 2022; Kang et al., 2014) (cluster HOP1), and those expressing prolactin-genes such as *TSC22D3, NDRG1* and *VEGFA* (Meng et al., 2019; Qiu, Bevan, et al., 2008; Sornapudi et al., 2018) (cluster HOP8), thus illustrating a degree of cell specificity in response to pregnancy hormones (**Fig. 4C-E**). We also identified differentiated population of LHS cells, marked by the expression of *CLND3* (clusters HOP5 and HOP6), and differentiated LHS cell cluster HOP9, which was biased towards conditions with shorter exposure to EPP, marked by the expression of *FXYD3* and *LCN2* (Y.-F. Liu et al., 2020; Xue et al., 2019) (**Fig. 4C-E**).

We also identified specific cytokeratin markers that further defined cellular states of human breast organoid cultures, with high levels of *KRT6* marking LASP and BMyo cells (clusters HOP1, HOP2, HOP4, and HOP8), and high levels of *KRT7* marking EPP-induced mature LHS cells (HOP5, HOP6, and HOP9) (**Fig. 4F**). Collectively, this analysis identified distinct cellular states, based on alterations to gene expression and organoid treatment response, thus illustrating the complex cellular dynamics induced by pregnancy hormones.

To further complement the molecular characterization of human breast organoid cultures, we employed a more general gene expression analysis, to indicate potential pathways enriched in breast epithelial organoid cultures (**Fig. S4D-G**). In doing so, we first examined enriched pathways of each cellular cluster from untreated human breast organoids. While luminal clusters defined to have a high proliferative state were marked by pathways associated with cell cycle regulation (clusters HOP3, HOP5), proliferating BMyo cluster HOP7 was marked by pathways linked with interferon responses, signals know to regulate the growth dynamics of epithelial cells (Cornelissen et al., 2020; Parker et al., 2016), thus suggesting distinct mechanisms of cell growth regulation in organoid systems (**Fig. S4D**). Interestingly, BMyo HOP2 cluster show no enrichment for a particular pathway, further suggesting an overall up-regulation of basal-like programs across populations of human breast organoid cultures (**Fig. S4D**).

Clusters of LHS cells were enriched with pathways associated with hormone response, such as TNF-⍺ signaling via NF-κB pathways, a pathway linked with increased mitogenic activity in response to estrogen (clusters HOP1 and HOP6), and mTOR signaling, which promotes proliferation in response to estrogen (clusters HOP4, HOP6 and HOP8) (**Fig. S4C**) (Ketterer et al., 2020; Morrison et al., 2015; Rubio et al., 2006). Cluster HOP4 and HOP6 were also enriched for genes associated with Hypoxia, thus suggesting cellular states with increased metabolic rates **(Fig. S4D**). Our analysis also identified LHS clusters enriched with hormone regulated pathways, such as Androgen response (cluster HOP6 and HOP9) and estrogen response (cluster HOP9), indicating highly hormone responding states in organoid cultures **(Fig. S4D**).

A similar analysis approach was employed to define the transcriptional state of cellular clusters in organoid cultures grown EPP (**Fig. S4E-F**). Our results suggest that while hormone clusters HOP5 and HOP6 show no enrichment for pathways when compared with no treatment, a metabolic state switch is suggested with prolonged pregnancy hormone exposure, with cluster HOP5 downregulating fatty acid associated signaling and HOP6 up-regulating process linked with p53 pathway, hypoxia and mTORC1 (**Fig. S4E-G**). Hypoxic-associated pathways were also identified in clusters HOP1 after 10 days of EPP culturing, and in clusters HOP8 and HOP9 across both EPP-treated conditions, further supporting the effects of pregnancy hormones on regulating the metabolic state of breast organoid cells (**Fig. S4E-G**). Cluster HOP3 was consistently enriched for genes linked with TNFa signaling via NF-Kb in response to EPP, while clusters HOP4 and HOP8 upregulated these signals only after prolonged EPP exposure (**Fig. S4D-G**).

Furthermore, cluster HOP4 was found to be enriched for additional pathways such as p53 pathways, mTORC signaling, glycolysis and androgen response after 21 days of EPP treatment (**Fig. S4G**). Prolonged exposure to pregnancy hormones also resulted on the enrichment of p53 pathway, oxidative phosphorylation in clusters HOP8 and HOP9, further demonstrating additional cell types with dynamic transcription regulation in response to hormones (**Fig. S4G**). Interestingly, cells from cluster HOP9 were also enriched for pathways associated with EMT and cMYC targets in response to prolonged exposure to EPP, thus suggesting the activation of cell plasticity process associated with pregnancy signals (**Fig. S4G**). Collectively, these findings illustrate the molecular and cellular alterations, induced by *ex vivo* exposure to pregnancy hormones, thus supporting the robustness of organoid cultures to understand normal developmental stages of human breast tissue.

### 5. Defining the evolutionary conserved responses to pregnancy hormones in mammary organoid cultures

Our analysis indicated that both murine and human mammary organoids treated with EPP recapitulated some of the previously described pregnancy-induced changes that take place *in vivo*. Yes, it is possible that pregnancy signals may activate pathways that are both evolutionary conserved and species specific. Therefore, we set out to define the evolutionary conserved basis of mammary organoid systems between human and murine cultures, by initially integrating untreated murine and human organoids datasets (referred hereafter as UMH clusters). Such approach identified a total of 7 clusters with varied distribution across species (**Fig. S5A**). To avoid lineage classification issues, biased by the state of human organoid cultures, we utilized once again the top differentially expressed genes to determine the identities of each UMH cluster (**Fig. S5B**).

Our analysis identified five clusters of luminal-biased cell types (UMH1, UMH3, UMH4, UMH5, and UMH7), from each two clusters were classified as LASP state (UMH1 and UMH4), and three clusters defined to be of LHS lineage (UMH3, UMH5, and UMH7), including two defined to be at high proliferative state (UMH3 and UMHM5). We also identified 2 clusters of BMyo cell types (UMH2, and UMH6), thus further supporting the heterogeneity of organoid derived from mammary tissue (**Fig. S5B-C**).

The distribution of organoid clusters also varied according to species. While clusters UHM1 (LASP) and UMH6 (BMyo) were biased towards samples from murine origin, BMyo (UMH2) and LASP (UMH4) fates were also identified in human organoid cultures, thus suggesting a species-specific distribution of these lineages in organoid cultures (**Fig. S5A and S5D**). Our analysis also identified clusters of cell populations somewhat present in both mouse and human organoid conditions, mostly represented by LHS lineages, including those at a high proliferating state (**Fig. S5A-D**). The aforementioned observations concur with previous findings comparing intact human and murine MECs, where luminal lineages, especially progenitor-like ones, were shared across species (Henry et al., 2021). Collectively, this approach allowed for the initial identification of species biased organoid cell types.

We next asked whether treatment with pregnancy hormones would influence the dynamics of species-specific mammary epithelial subtypes. In doing so, we integrated EPP-treated murine and human organoids datasets (referred hereafter as PMH clusters), an approach that yield 9 cellular clusters of several epithelial lineages (**Fig. 5A-B**). Our analysis once again identified cell populations that are shared across species, and those that are species specific, thus illustrating differences to how breast organoid cultures from mouse and human mammary glands respond to pregnancy hormones (**Fig. C-F**).

**Figure 5.**
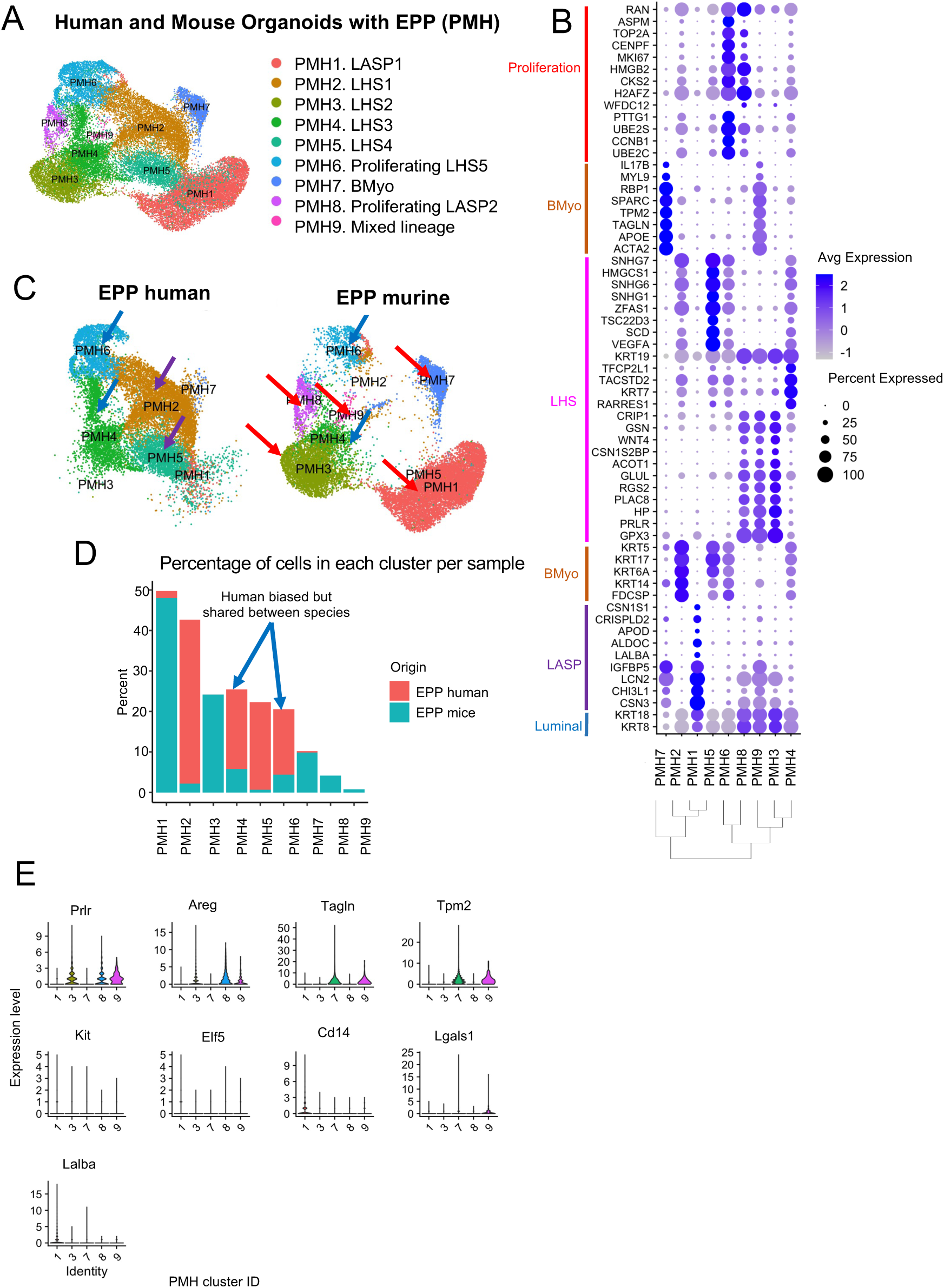
Evolutionary comparisons between EPP-treated murine and human organoid MECs. (A) Resulting clusters of human and mouse organoids with EPP (PMH). PMH clusters were assigned their respective identities based on previously described human MEC markers gene expression and their top DEGs. (B) Dotplot showing expression of the top DEGs per PMH cluster. Clusters are organized based on their dendrogram relationships. (C) PMH clusters split by species of origin. The purple arrows highlight clusters enriched in human samples, red arrows highlight clusters enriched in mouse samples, and blue arrows highlight clusters enriched in both species. (D) Bar plot showing percentage of cells per condition in each PMH cluster. (E) Violin plots for expression of genes characterizing mouse-biased clusters from PMH. Clusters enriched in both humans and mice are highlighted by the blue arrow.

For example, BMyo cell types (PMH7) and mixed lineage cell types (PMH9) were identified to be biased to mouse organoids treated with EPP, perhaps illustrating distinct alterations to cellular states in response to pregnancy hormones compared to human organoids (**Fig. 5B-D**). In addition, we identified several clusters of cells bearing high levels of *CSN3* mRNA, a gene that is associated with a LASP state in clusters that were either made up of mouse and human MECs (cluster PMH1) or in clusters biased towards mouse organoid conditions that also expressed elevated *PRLR* mRNA levels (clusters PMH3, PMH8 and PHM9) (**Fig. 5B-C** **and 5E**). These observations suggest that prolactin-induced responses are more efficiently activated in murine systems supplemented with EPP.

Human organoid biased clusters were identified to bear an LHS state (cluster PMH5), a cell type showed to be expanded by EPP, thus indicating selective responses by organoid treatment (**Fig. 4 and 5B-C**). In fact, additional clusters of cells with representation in both murine and human datasets were classified to have an LHS lineage, including those in highly proliferative states, that were induced by EPP treatment, (**Fig. 5B-C, and S5E**). Therefore, our findings support that our organoid conditions, to some extent, recapitulates pregnancy-induced development observed in both mouse and human mammary systems (**Fig. 5B-E**).

## Discussion

Our characterization of MEC-derived organoids at a single-cell level allowed us to carry out a comprehensive assessment of organoid systems to model mammary gland development. Our initial analysis of murine MEC-derived organoids scRNA-seq data confirmed conservation of *in vivo* lineage signatures, as well as representation of a diverse array of MEC lineages *ex vivo*. These results complement a previous proteomics study that made use of Cytometry by time of flight (CyTOF) to confirm that MEC lineages found *in vivo* are present in patient MEC-derived organoid cultures (Gray et al., 2022). We further confirmed lineage fidelity between *in vivo* and 3D *ex vivo* systems by comparing scRNA-seq data from intact murine mammary tissue to data we generated from murine MEC-derived organoids. This particular analysis resulted in the appearance of a luminal progenitor population that is organoid exclusive, suggesting that certain cells in culture exist in a stem-like state, perhaps responsible for maintaining the growth of organoids *ex vivo*. Although organoids successfully replicate the composition of primary tissue, we also observed exclusive organoid states in both non-stimulated and hormone-stimulated conditions, indicating that discrepancies between the systems may contribute to the emergence of these phenotypes in culture.

The induction of MECs into an immature cellular state in organoid cultures could have resulted from a lack of microenvironment queues that are crucial for mammary development. For example, prior research has highlighted the significance of various fibroblast types in MEC development and homeostasis, as well as the potential role of adipocytes in regulating MEC growth and function stages (Gregor et al., 2013; Hovey & Aimo, 2010; Howard & Lu, 2014; X. Liu et al., 2012; Makarem et al., 2013; X. Wang & Kaplan, 2012). Moreover, signals that are not necessarily produced by surrounding non-epithelial cells in the mammary gland but that can result from paracrine signaling from other tissues are also vital for the maturation of specific MECs, such as OT, which promotes the differentiation of myoepithelial cells (Sapino et al., 1993). Furthermore, media composition has been shown to affect organoid culture composition (Gray et al., 2022), which could also have contributed to the observed phenotype. Nonetheless, regulon analysis revealed that regulons with a high RSS for each system overall contributed to the needs of each system to survive and reach a homeostatic state in their respective microenvironments. Thus, MEC-derived organoids are a suitable system to assess the effect of controlled developmental signals, but should be used with the previously discussed considerations. Future studies involving the addition of signals that contribute to endogenous mammary gland development and maintenance, along with co-culturing with essential cells from the mammary microenvironment will further improve the fidelity of this system (Koledova & Lu, 2017; Sumbal et al., 2021, 2022). Moreover, assessment of secreted factors by these non-epithelial cells and estimation of direct cell-cell communication dynamics would provide valuable insights. Additional signals known to promote MEC maintenance and development could be introduced, such as the addition of Oxytocin to the cultures. The addition of Oxytocin has the potential to promote pseudo-lactation, shifting the organoid cultures towards a lactation-like heterogeneity rather than a pseudo-gestation state (Sumbal et al., 2020).

Single-cell characterization of murine MEC-derived organoids treated with different concentrations of estrogen enabled us to begin to isolate the effects of individual hormones on MEC development, especially during distinct biological processes involving an interplay of varying hormone doses (e.g. the estrus cycle). This analysis revealed the emergence of an estrogen-exclusive LHS population independent of dosage, as well as a depletion of mixed lineage cells exclusively at a higher dose of estrogen. Our results suggest that LHS cells in our estrogen-exclusive cluster are potentially cells that were already present in organoids without treatment, in a cellular state triggered by hormone treatment. This is evidenced by the simultaneous depletion of a cellular cluster of LHS cells enriched in untreated samples. Further comparison of both LHS clusters revealed that estrogen-exclusive LHS cells highly express *Areg* and *Pgr*, both which have been previously described to be upregulated by estrogen (Kanaya et al., 2019). Moreover, our findings that estrogen-exclusive LHS cells are highly differentiated compared to untreated LHS cells indicate that hormone treatment could be promoting cellular maturation, in accordance with previous studies indicating that a LHS cell mature state is correlated with expression of hormone receptors (Bach et al., 2017). These mature LHS cells also displayed an activation of pathways associated with proliferation and inflammation, which have been previously linked to estrogen-mediated activation (Maharjan et al., 2021). Conversely, in mixed lineage cells, an activation of genes associated with Androgen response was observed. Androgen receptors can block the pro-proliferative role of estrogen, suggesting a mechanism by which these cells are depleted in the presence of a certain estrogen dose (Bleach & McIlroy, 2018).

A lack of hormone signals at baseline could further explain why we observe an enrichment of mixed lineage cells in organoids without treatment and a stark depletion in organoids treated with a high dose of estrogen. This interpretation is complementary to a previous study that delineates a quiescent state for mixed lineage cells in the adult mammary gland, which become active in the presence of hormones (Fu et al., 2017).). Therefore, these results highlight that organoid culturing conditions at baseline resemble developmental stages found in a microenvironment depleted of hormones, such as prepubescent development and menopause. Given that an aged extracellular matrix alone can drive MECs into neoplastic and invasive cellular states (Bahcecioglu et al., 2021), it will be important to identify what stages of development the composition of Matrigel and organoid media resembles most. Thus, our analysis paves the way to future studies that will involve comparing organoid MECs with intact MECs from pre-pubescent and post-menopausal mice, as well as more studies involving dissecting the individual effects of hormones on MEC development and maintenance.

Previous work using a combination of prolactin, hydrocortisone, OT, and growth factors showed mouse MEC-derived organoids are able to mimic lactation and involution (Sumbal et al., 2020). Additional studies further introduced the idea of using a cocktail of pregnancy hormones (estrogen, progesterone and prolactin, or EPP) without growth factors to simulate a pseudo-lactation state, which resulted in the incremental expression of *Csn2* and changes to the epigenome previously associated with pregnancy (Ciccone et al., 2020). Our current study extends upon these studies by demonstrating compositional and transcriptomic changes to mammary organoids as a direct effect of treatment with pregnancy hormones. We show a depletion and emergence of similar cell types with pregnancy hormones treatment, suggesting that the observed compositional differences between organoids without treatment and with pregnancy hormones are likely due to subtle changes in cellular states. Moreover, cellular clusters that emerge with pregnancy hormones treatment are enriched for processes that have been previously associated with lactation, such as adipogenesis and hypoxia (Colleluori et al., 2021; Y. Shao & Zhao, 2014). Therefore, these results indicate specific cell types obtain a parity-associated gene expression signature with exposure to hormones during pregnancy. We further compared scRNA-seq data from MECs obtained at intact pregnancy stages (Bach et al., 2017) with our organoids treated with pregnancy hormones, and found our organoid cultures recapitulate lineages from all pregnancy stages.

We also found that our organoids with and without pregnancy hormones treatment both have cellular populations present in a cellular state only found in MECs undergoing gestation, thus suggesting that the proliferative and stem-like state of organoid MECs is most similar to this stage of pregnancy. Additionally, the emergence of a cellular state exclusive to organoid cultures treated with pregnancy hormones once more suggests that certain phenotypes are exclusive to our culturing conditions, even in the presence of hormones. Therefore, we conclude that organoids can recapitulate drastic cellular changes that occur with pregnancy, particularly by mimicking the gene signature of MECs during pregnancy. However, since organoid MECs at baseline appear to have additional levels of proliferation than nulliparous MECs, this model must be used with caution to understand other aspects of pregnancy-associated development.

In fact, analysis comparing untreated and treated organoid cultures identified a population of LHS MECs largely exclusive to conditions supplemented with pregnancy hormones, thus supporting a possible cellular expansion in response to pregnancy signals (**Fig. 3A-B**). Interestingly, the existence of pregnancy-induced MECs (PI-MECs) has already been suggested in intact mammary tissue, although its true lineage identity and function remain very controversial (Chang et al., 2014). Such pregnancy-induced, stably sustained cellular state, could also represent populations that bear pregnancy-induced epigenetic changes, and therefore the cellular basis for a robust response to consecutive exposures to pregnancy signals (Ciccone et al., 2020; dos Santos et al., 2015; Feigman et al., 2020).

We were able to uncover the translational potential of MEC-derived organoids by further showing that patient MEC-derived organoids respond to pregnancy hormones by inducing transcriptomic changes to organoid MECs associated with pregnancy. Interestingly, similar to our observation in murine organoids, one of the enriched pathways in human organoids treated with pregnancy hormones were those associated with hypoxia, reflecting its significance in pregnancy where the mammary gland boosts metabolic activity to support growth and lactogenesis, thereby activating hypoxia-associated genes (Y. Shao & Zhao, 2014). Particularly, hypoxia stabilizes HIF-1α from the HIF complex, which upregulates glucose intake in the mammary gland. Interestingly, we observed a coupled enrichment of glycolysis-associated genes in human organoids treated with pregnancy hormones, potentially indicating a striking role of hypoxia in preparing mammary organoids for pseudo-lactation.

Nonetheless, we found that most human organoid MECs exist in a luminal-basal state. The phenomenon of organoids becoming more basal-like after long term culturing had already previously been reported (Bhatia et al., 2022), thus potentially confirming that the phenotype we observed in human organoids could be a result of the number of passages prior and during the course of the experiment. One approach that could be implemented to address this issue is to grow the cells and sequence them right before the next passaging, allowing cells to differentiate in culture prior to sequencing. However, there are other factors that could affect the observed phenotypes in culture, such as the inability to remove growth factors from culture due to the developmental timeline of human organoid MECs compared to murine organoids. Notably, despite the mixed lineage phenotype we observed, we did identify LHS cells with low levels of ERα expression and expression of downstream estrogen targets in these human MEC-derived organoid cultures. It has long been a challenge to obtain hormone positive clones in culture, as previous studies using BC-derived organoids have noted that the expression of hormone receptors is reduced in culture compared to intact tissue (Campaner et al., 2020; Guillen et al., 2022). Our identification of LHS clones in normal MEC-derived organoids using both a combination of hormone receptor status and downstream targets therefore suggests the potential of using these 3D cultures for understanding the development of hormone positive BCs, extending the applications of 3D cultures towards both fundamental biological research and potential clinical implications.

In the comparison of untreated organoids between human and mouse, mature cells clustered mainly in a species-specific manner, preserving the suggested hierarchy across species while displaying divergent epithelial responses, as it has been reported in previous literature (Henry et al., 2021). Interestingly, a subset of proliferative LHS cells was identified in both human and murine MEC organoid cultures, suggesting a conserved population that plays a crucial role in maintaining mammary tissue homeostasis throughout evolution. However, upon treatment with pregnancy hormones, further significant differences in mature cell types emerged between the species. The compositional differences observed in milk from various mammalian species, influenced largely by phylogeny, imply intrinsic cellular response variations (Skibiel et al., 2013). Notably, the shared LHS cell population between mice and humans in cultures treated with pregnancy hormones appeared to be divided between proliferating and non-proliferating cells, indicating potential expansion and distinct functions in preparing the mammary gland for lactation. Moreover, shared cluster signatures across species in response to pregnancy hormones highlighted processes associated with pregnancy, such as hypoxia, estrogen response, and fatty acid metabolism (Colleluori et al., 2021; Y. Shao & Zhao, 2014). Therefore, our findings shed light on the intricate interplay between species-specific and conserved cellular responses in the context of mammary tissue dynamics and lactation preparation. Furthermore, our analysis demonstrates the utility for comparing epithelial responses of organoids cultured from different species in order to identify evolutionarily conserved cellular states that could be highly relevant for mammary tissue homeostasis during different developmental processes.

Altogether, we have developed an atlas of normal MEC-derived organoids from mouse and human tissue, which can be incorporated with other single-cell methods to understand the molecular mechanisms governing MEC development *ex vivo*. We characterize the effects of feminizing hormones on these 3D cultures at a single-cell level, supporting hormone treatment of organoids as a system to understand developmental processes associated with adolescence, pregnancy and menopause. Our findings support the implementation of this procedure as a non-invasive method to understand how the human mammary gland is modified during a pregnancy cycle. This system can also be extended to other species, in order to assess the evolutionary basis of MEC response to hormones across other mammalian species.

## Data availability statement

scRNA-seq datasets were deposited into NCBI database [https://www.ncbi.nlm.nih.gov/], BioProject PRJNA1015687, and will be made publicly available upon manuscript publication. Previously published datasets are available under the following IDs: SAMN16776241 (re-clustered MECs from nulliparous BALB/c mice), GSM2834498, GSM2834499 (mammary tissue from nulliparous C57BL/6 female mice), GSM2834500, GSM2834501 (mammary tissue from C57BL/6 female mice at mid-gestation), GSM2834502, GSM2834503 (mammary tissue from C57BL/6 female mice during lactation), GSM2834504, GSM2834505 (mammary tissue from nulliparous C57BL/6 female mice at late involution stage).

## Author contributions

C.O.D.S. oversaw the project, and participated in experimental design, data analyses and manuscript preparation. S.M.L. and M.F.C. cultured the organoids and prepared cells for single-cell RNA-seq libraries. J.R.O. performed quality control steps on resulting scRNA-seq files, as well as all subsequent bioinformatics analysis steps. S.H. provided code for data integration, and D.C. set up the SCENIC runs and provided initial code for subsequent regulon analyses. A. S. provided vital feedback for the evolutionary analyses performed in this project.

## Acknowledgements

This work was performed with assistance from CSHL Animal Facility, the CSHL NextGen Sequencing Shared Resources and the CSHL Single Cell Shared Resources, which are supported by the CSHL Cancer Center Support Grant 5P30CA045508. This work was financially supported by the CSHL and Northwell Health affiliation, the NIH/NCI grant 1R01CA248158-01, the NIH/NIA grant R01AG069727-01, and the NIH/NCI 1R01CA284630 (C.O.D.S.). We would also like to acknowledge Asma Kaleem for helping with the culturing of human MEC-derived organoids, and Dr. Yixin Zhao for sharing initial code for evolutionary comparisons.

## Supplementary Figures

**Supplementary Figure 1.**
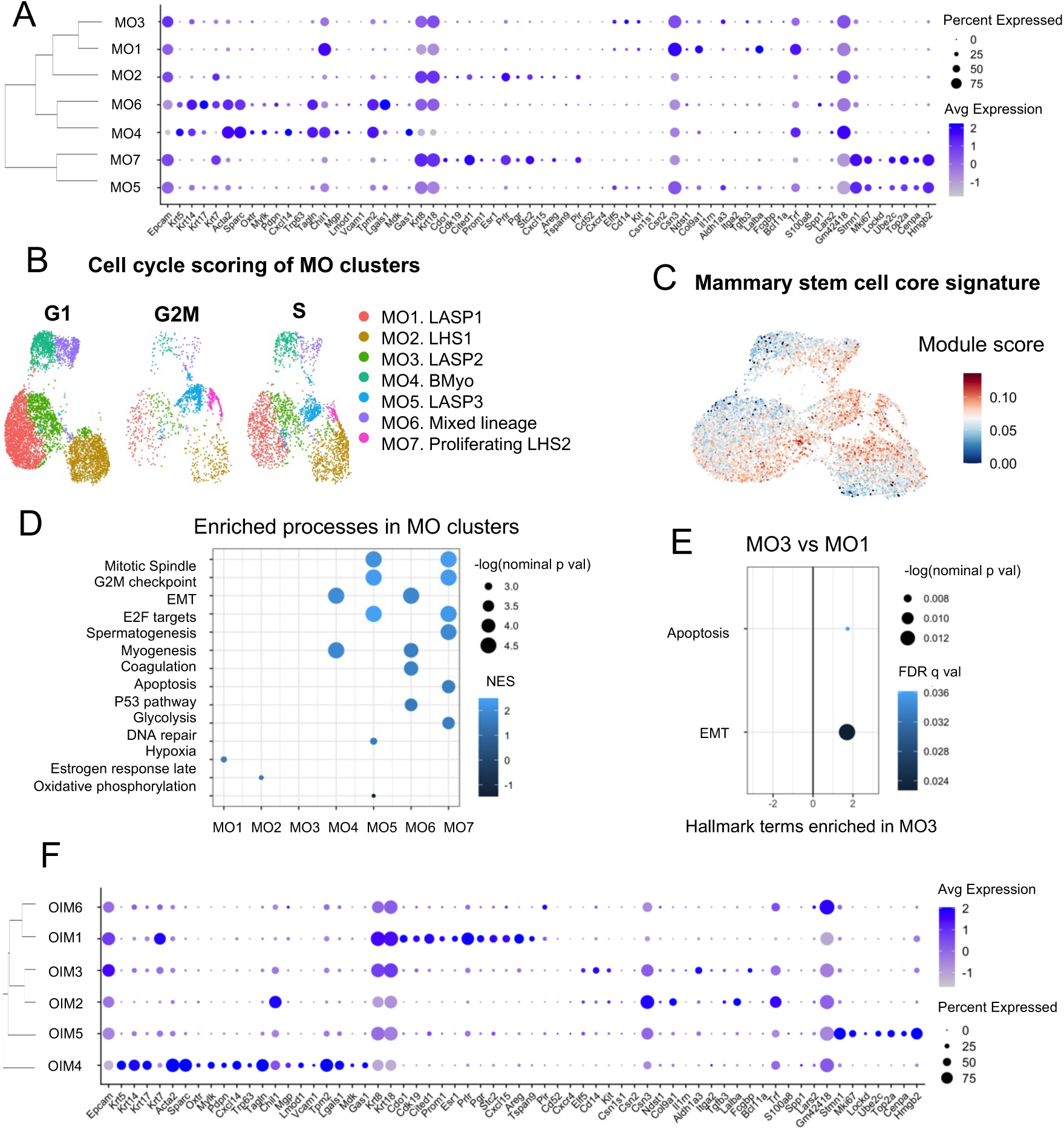
Approaches for MO and OIM characterizations. (A) Dotplot showing MEC markers average gene expression in each MO cluster. (B) Cell cycle scoring of MO clusters. (C) MO clusters scores for mammary stem cell core signature retrieved from Enrichr. Dark blue represents a score of 0 and red indicates a score of 0.10. (D) Summary of enriched hallmark terms in each MO cluster, ordered based on each –log(nom p-val) for each term. Only terms with nom p-val < 0.05 were kept for this analysis. The color of each dot represents the NES value for each term. (E) GSEA for enriched hallmark terms in cluster MO3 vs MO1. The terms are ordered based on each –log(nom p-val) for each term. Resulting terms were not filtered for significance given that none passed the p-val < 0.05 threshold. (F) Dotplot for MEC marker expression from OIM clusters. OIM clusters are organized based on dendrogram relationships.

**Supplementary Figure 2.**
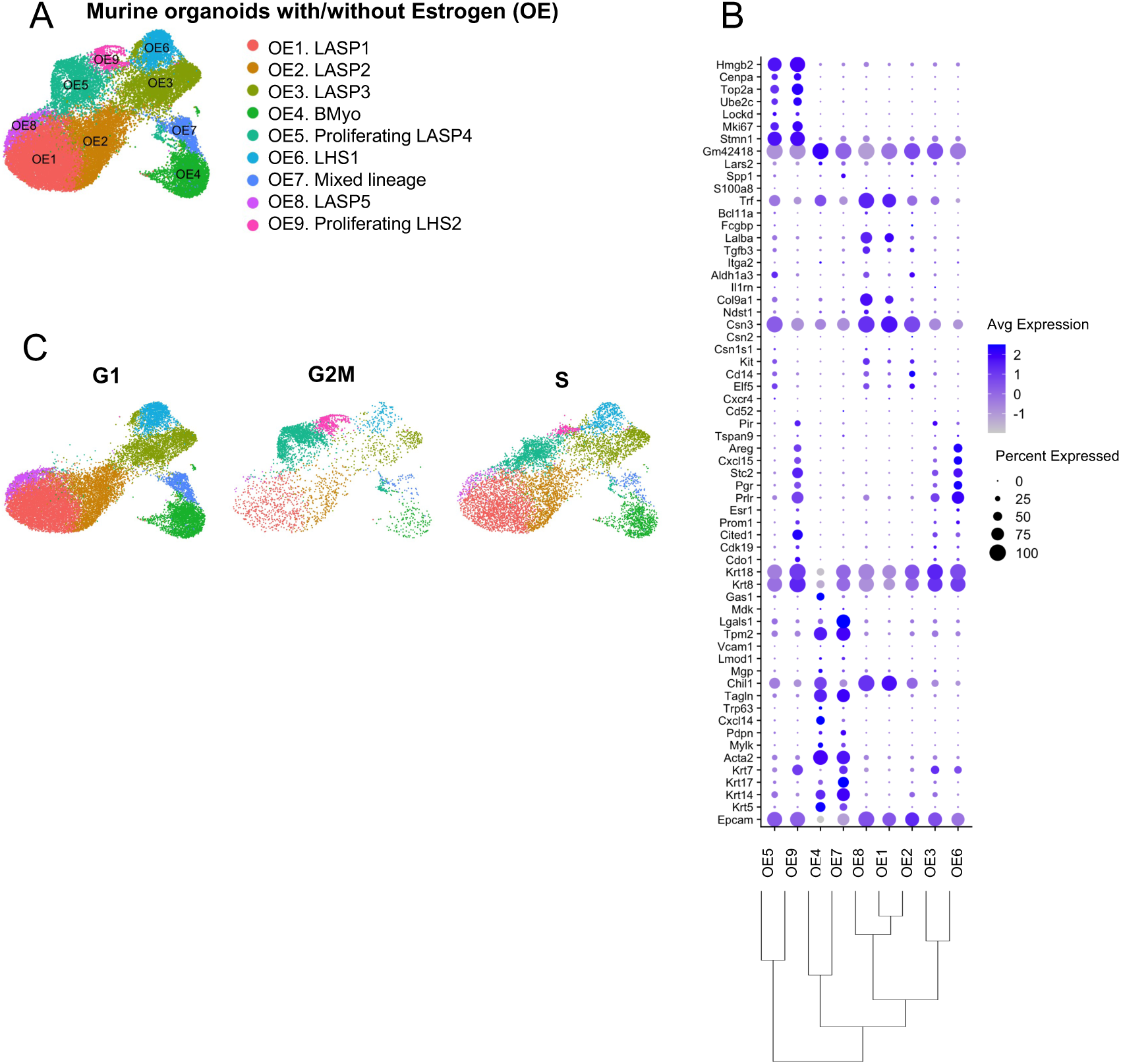
Strategies for classifying OE clusters. (A) Resulting clusters for murine organoids with and without estrogen (OE), with cluster identities based on intact MEC marker expression. (B) Dotplot for OE clusters expression of intact MEC gene markers. OE clusters are organized based on dendrogram relationships. (C) Cell cycle scoring or OE clusters.

**Supplementary Figure 3.**
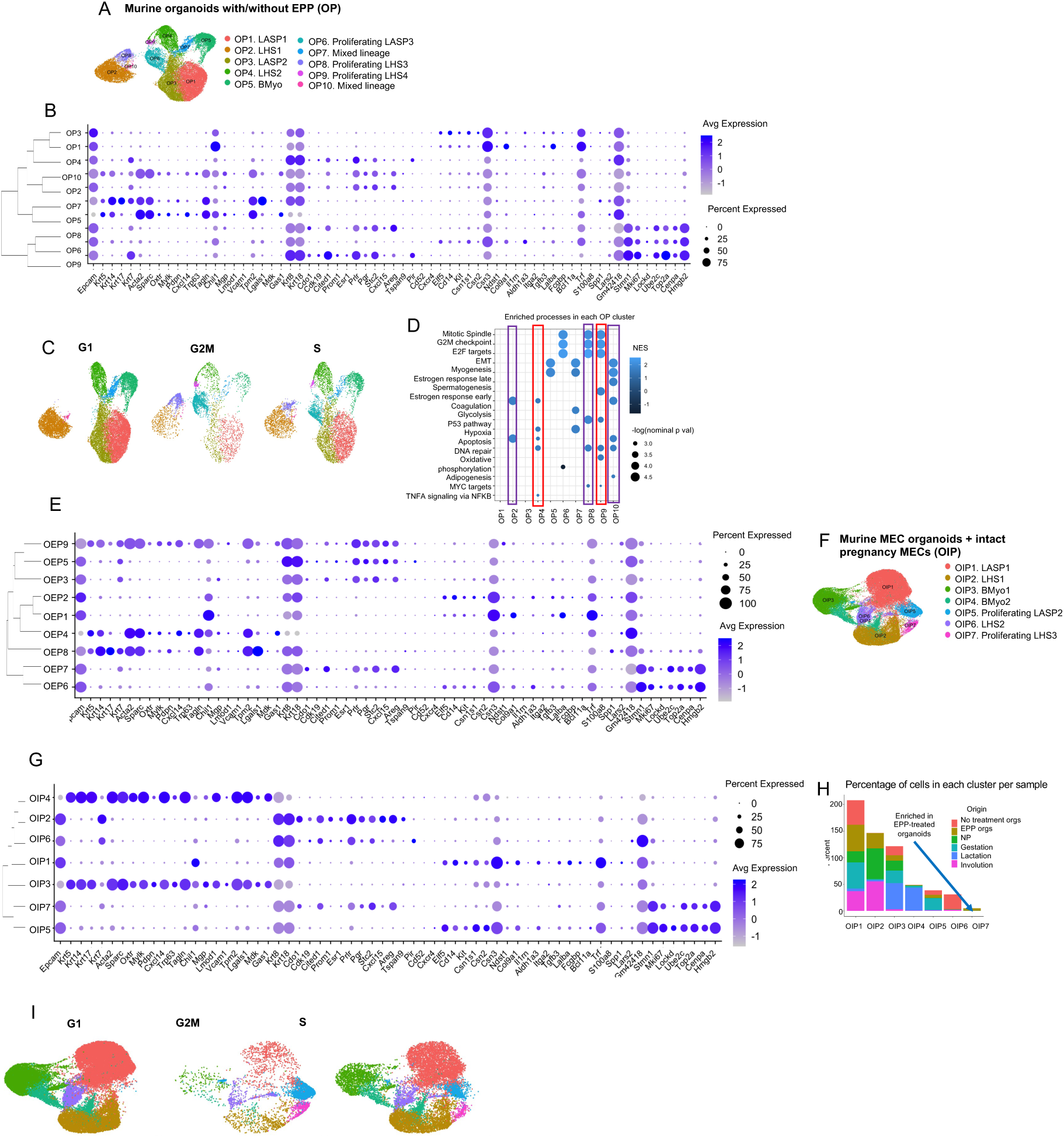
Classification strategies for OP and OIP clusters. (A) Resulting clusters of murine organoids with and without EPP (OP), with their identifications corresponding to expression of previously known MEC markers. (B) Dotplot for MEC marker expression in OP clusters. Clusters are organized based on dendrogram relationships. (C) Cell cycle scoring of OP clusters. (D) GSEA for hallmark terms enriched in each OP cluster. Terms are ordered from highest –log(nom p-value). Only hallmark terms with a normalized p-val < 0.05 were kept for this analysis. The color of each dot represents the NES for each term. The red boxes mark clusters depleted with EPP treatment, and the purple boxes mark clusters enriched with EPP. (E) Dotplot for MEC marker expression in OEP clusters. Clusters are organized based on dendrogram relationships. (F) Resulting clusters of murine organoids with and without EPP treatment integrated with data sets from intact pregnancy MECs (OIP), with their identifications corresponding to expression of previously known MEC markers. (G) Dotplot for MEC marker expression in OIP clusters. Clusters are organized based on dendrogram relationships. (H) Bar plot showing percentage of cells per condition in each OIP cluster. The blue arrow highlights cluster OIP7, which is enriched in EPP-treated organoids. (I) Cell cycle scoring of OIP clusters.

**Supplementary Figure 4.**
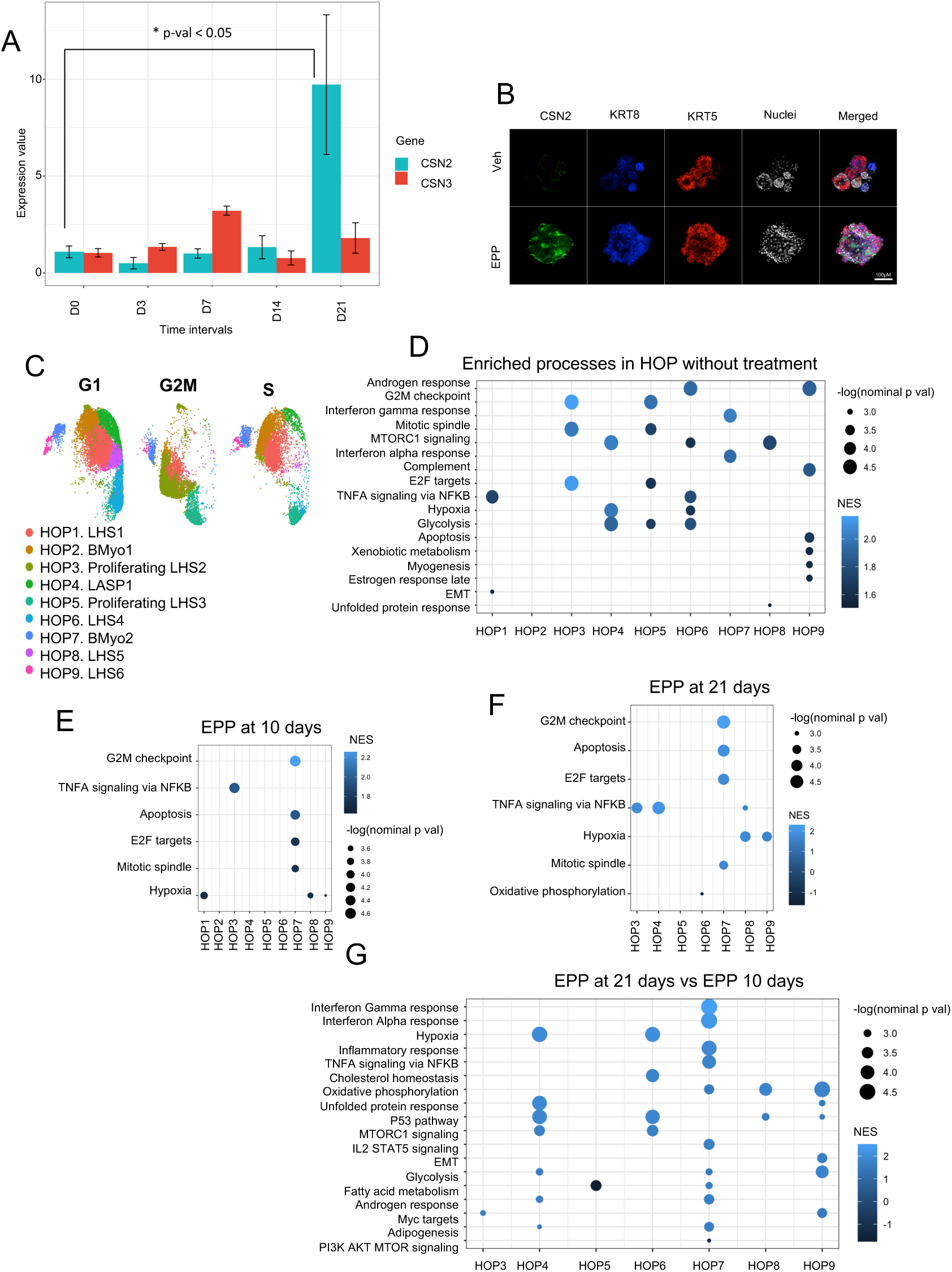
Strategies for determining the optimal timeline and treatment effects of EPP on human organoids through gene expression analysis. (A) qPCR results for CSN2/CSN3 expression in human MEC-derived organoids treated with EPP for 21 days (n=2 runs per day). Comparing CSN2 expression at 0 days and 21 days of EPP treatment resulted in a significant difference in expression (p-val = 0.0238). (B) Immunofluorescence stain for milk-associated protein CSN2 in human organoids treated with pregnancy hormones (EPP). (C) Cell cycle scoring of each HOP cluster. (D) GSEA for enriched hallmark terms in each HOP cluster with no EPP treatment. Terms are ordered based on–log(nom p-value). Only hallmark terms with a normalized p-val < 0.05 were kept for this analysis. The color of the dots represents NES. (E-G) GSEA for hallmark terms differentially enriched in EPP-time point across clusters. Terms were ordered decreasingly based on –log(nom p-value). Only terms with nom p-val <0.05 were kept for these analyses. Likewise, the color of the dots represents NES.

**Supplementary Figure 5.**
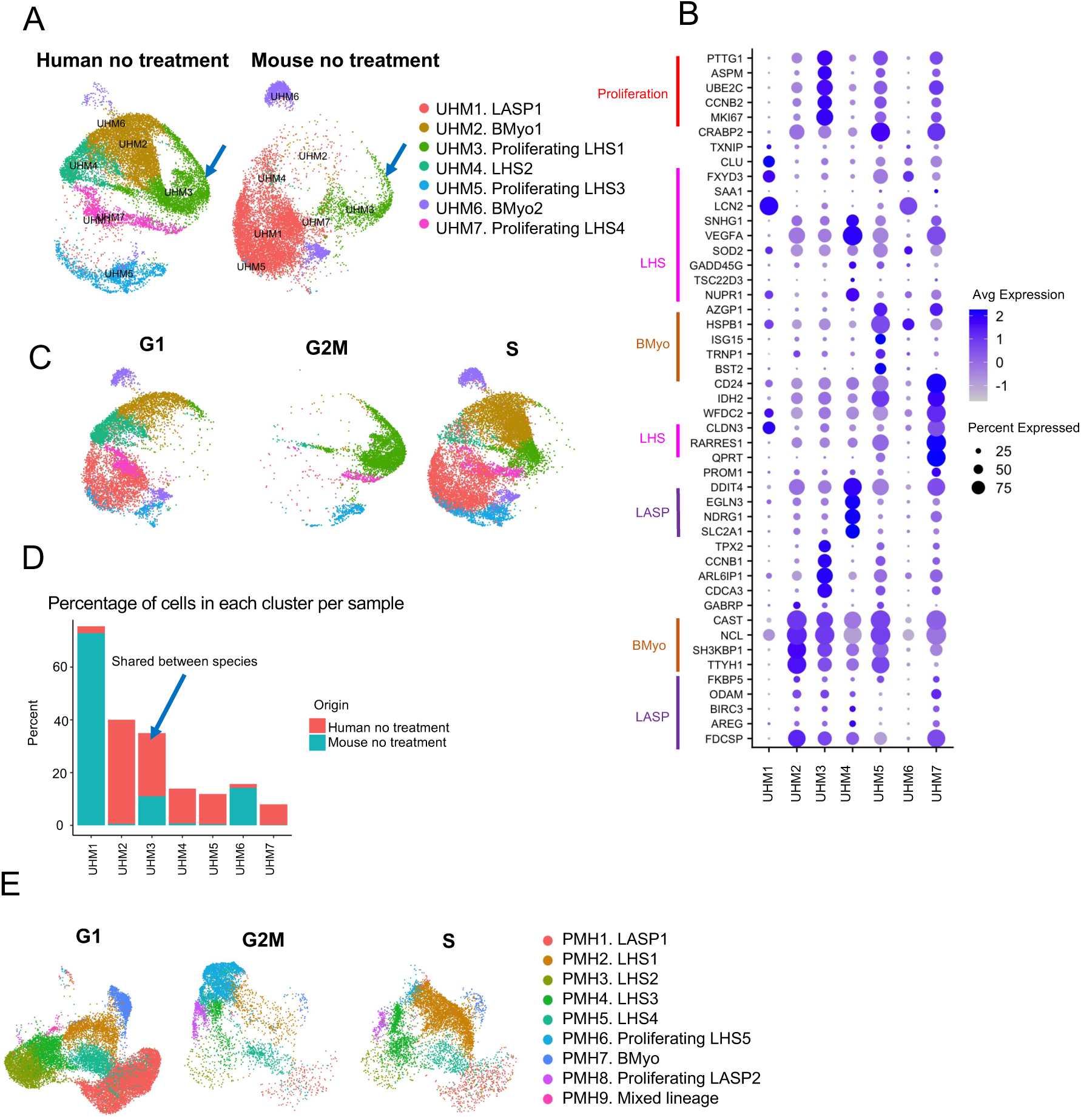
Additional transcriptomic analyses for MHP clusters. (A) Integration of data sets from murine and human organoid MECs without treatment (Untreated and Human MEC organoids – UHM). The blue arrow highlights a cellular cluster shared between species. (B) Dotplot showing expression of the top DEGs per UHM cluster. (C) Cell cycle scoring of UHM clusters. (D) Bar plot showing percentage of cells per condition in each UHM cluster. Clusters enriched in both humans and mice are highlighted by the blue arrow. (E) Cell cycle scoring of PHM clusters.

## Supplementary Tables

**Table S1.**
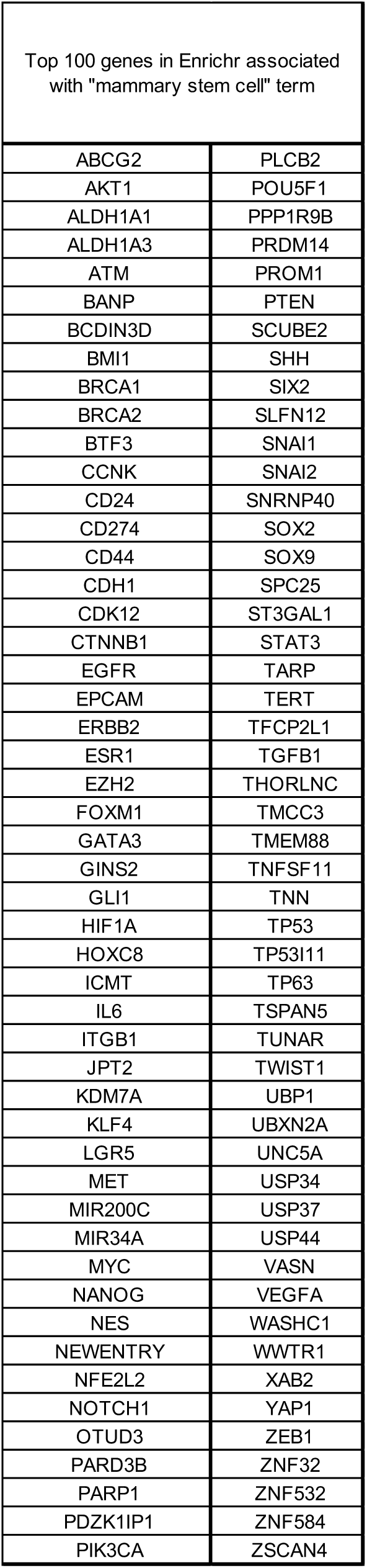
Top 100 genes in Enrichr associated with “mammary stem cell” term.

**Table S2.** GSEA terms associated with hypoxia and coagulation enriched in MO clusters.

| HALLMARK TERM | CLUSTER | GENE | RUNNING ES | RANK IN GENE LIST | RANK METRIC SCORE |
| --- | --- | --- | --- | --- | --- |
| HYPOXIA | MO1 | LALBA | 0.1767235 | 0 | 2.702094793 |
|  |  | ALDOC | 0.314538 | 1 | 2.107177734 |
|  |  | AKAP12 | 0.3647591 | 13 | 1.38170886 |
|  |  | CITED2 | 0.38943002 | 26 | 1.046850681 |
|  |  | HSPA5 | 0.39816093 | 40 | 0.858931601 |
|  |  | ERRFI1 | 0.3969774 | 55 | 0.763142109 |
|  |  | IER3 | 0.39405325 | 69 | 0.680725873 |
|  |  | ALDOA | 0.4308381 | 72 | 0.674044549 |
|  |  | TGFB3 | 0.43979836 | 82 | 0.639226735 |
|  |  | PLIN2 | 0.4464105 | 92 | 0.603324473 |
|  |  | HK1 | 0.47339243 | 96 | 0.579960883 |
|  |  | HS3ST1 | 0.5072853 | 98 | 0.574023366 |
|  |  | PFKL | 0.5060215 | 109 | 0.538704038 |
|  |  | ZFP36 | 0.51427794 | 117 | 0.516859949 |
|  |  | ENO1 | 0.47632548 | 137 | 0.479961067 |
|  |  | BCL2 | 0.45748258 | 151 | 0.437328964 |
|  |  | CCN1 | 0.40651336 | 173 | 0.392540962 |
|  |  | FOS | 0.27737224 | 215 | 0.313350797 |
| COAGULATION | MO6 | FN1 | 0.19547978 | 0 | 3.089641333 |
|  |  | SPARC | 0.30534342 | 11 | 2.065721273 |
|  |  | KLK8 | 0.34564635 | 33 | 1.328492403 |
|  |  | PRSS23 | 0.4036415 | 42 | 1.180061698 |
|  |  | BMP1 | 0.41370735 | 66 | 0.916438878 |
|  |  | HTRA1 | 0.44462416 | 78 | 0.850861132 |
|  |  | PDGFB | 0.49842855 | 79 | 0.850401461 |
|  |  | CSRP1 | 0.5167443 | 95 | 0.783407629 |
|  |  | THBS1 | 0.5432527 | 106 | 0.748256207 |
|  |  | ANXA1 | 0.5701388 | 116 | 0.721297979 |
|  |  | TIMP3 | 0.58145714 | 132 | 0.672811389 |
|  |  | LRP1 | 0.4589744 | 208 | 0.533705413 |
|  |  | TIMP1 | 0.46154517 | 223 | 0.501624227 |
|  |  | MMP14 | 0.32360157 | 302 | 0.388124615 |
|  |  | ITGB3 | 0.337482 | 308 | 0.384025753 |
|  |  | WDR1 | 0.07166303 | 445 | 0.27682212 |
|  |  | CAPN2 | 0.01946802 | 479 | 0.261658788 |
|  |  | KLF7 | 0.004166544 | 495 | 0.252073228 |

**Table S3.**
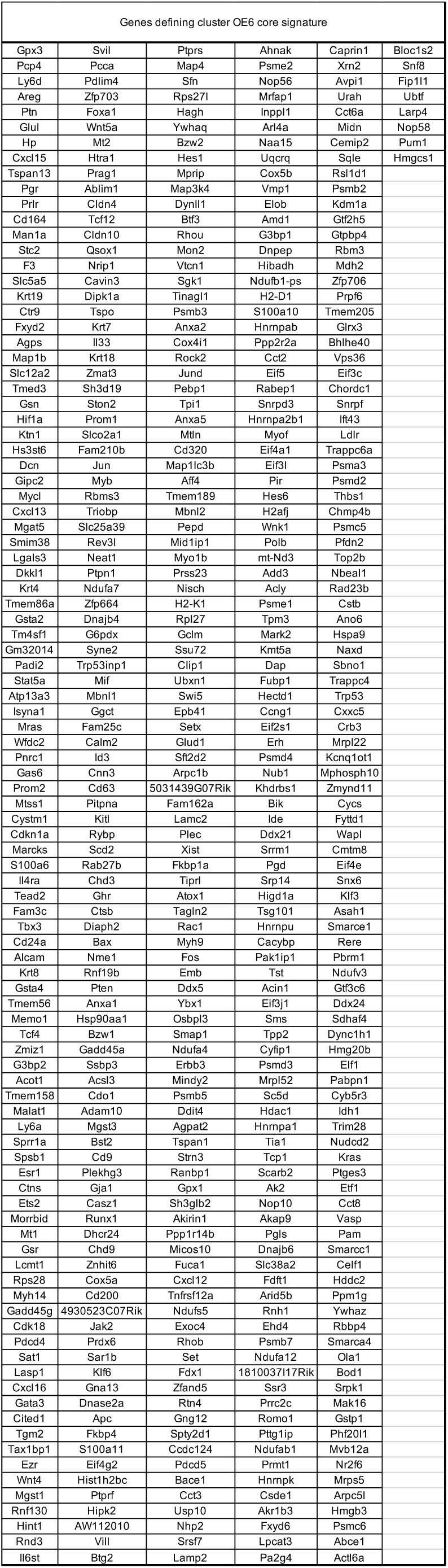
Genes defining cluster OE6 core signature.

